# Isoform-specific regulation of rhythmic gene expression by alternative polyadenylation

**DOI:** 10.1101/2020.12.12.422514

**Authors:** Ben J Greenwell, Joshua R Beytebiere, Teresa M Lamb, Deborah Bell-Pedersen, Christine Merlin, Jerome S Menet

## Abstract

Alternative polyadenylation (APA) generates transcript isoforms with different 3’ ends. Differences in polyadenylation sites usage, which have been associated with diseases like cancer, regulate mRNA stability, subcellular localization, and translation. By characterizing APA across the 24-hour day in mouse liver, here we show that rhythmic gene expression occurs largely in an APA isoform-specific manner, and that hundreds of arrhythmically expressed genes surprisingly exhibit a rhythmic APA isoform. The underlying mechanisms comprise isoform-specific post-transcriptional regulation, transcription factor driven expression of specific isoform, co-transcriptional recruitment of RNA binding proteins that regulate mRNA cleavage and polyadenylation, and, to a lesser extent, cell subtype-specific expression. Remarkably, rhythmic expression of specific APA isoforms generates 24-hour rhythms in 3’ UTR length, with shorter UTRs in anticipation of the mouse active phase. Taken together, our findings demonstrate that cycling transcriptomes are regulated by APA, and suggest that APA strongly impacts the rhythmic regulation of biological functions.

## Introduction

Most genes in higher eukaryotes contain multiple sites at which RNA can be cleaved and polyadenylated, leading to the expression of several distinct transcript isoforms (Reyes and Huber, 2018; Wang et al., 2008). The usage of these alternative polyadenylation (APA) sites affects 3’ UTR length and/or protein sequence, and leads to transcripts being truncated or containing additional exonic or intronic sequences (Tian and Manley, 2013, 2017). The length of the 3’ UTR tail itself has large implications on RNA stability, mostly because it defines the probability for a 3’ UTR to be targeted by RNA binding proteins (RBPs), miRNA, or long non-coding RNA (Gong and Maquat, 2011; Licatalosi et al., 2008; Sandberg et al., 2008). 3’ UTR length also affects the nuclear export and cytoplasmic subcellular localization of many mRNAs, thus allowing for the enrichment of proteins at a specific cellular location (An et al., 2008; Martin and Ephrussi, 2009). Differences in mRNA 3’ UTR length can additionally lead to differences in the localization of the resulting protein independently of mRNA localization, suggesting that 3’ UTRs can also bear crucial roles in the function of the protein they encode (Berkovits and Mayr, 2015).

Investigation of gene expression at the genome-wide level is commonly achieved through the use of massively parallel RNA-Seq libraries, where analysis frequently quantifies expression across all potential isoforms and reports a single value for each gene. Thus, RNA-Seq analysis commonly relies on the assumption that all distinct isoforms of each gene are regulated in a similar manner. However, it remains unclear whether APA isoforms are for the most part regulated in a similar manner, or if differential regulation of APA isoform expression occurs, and, if so, how this may affect downstream biological functions. Recent evidence has shown that biological processes like pluripotency are regulated by APA (Modic et al., 2019; Sandberg et al., 2008; Ye and Blelloch, 2014), and that defects in APA can lead to health defects including cancer (Gruber and Zavolan, 2019; Mayr and Bartel, 2009; Rehfeld et al., 2013; Weng et al., 2019). The underlying mechanisms involve, at least in part, the co-transcriptional loading of RBPs, which promote cleavage and polyadenylation at proximal APA sites to lead to increased expression of APA isoforms with shorter 3’ UTRs and decreased expression of isoforms with long 3’ UTRs. Thus, APA seem to mostly involve changes in the relative ratio of short *vs*. long APA isoforms without affecting the overall level of transcription initiation (Gruber and Zavolan, 2019; Xu and Zhang, 2018).

Regulation of rhythmic gene expression by the circadian clock has been described in numerous species and tissues, and about half of the transcriptome in mammals is rhythmically expressed in at least one tissue (Mure et al., 2018; Ruben et al., 2018; Zhang et al., 2014). This widespread rhythmic expression is generated by a transcriptional/translational negative feedback loop, which in mammals is initiated by the heterodimeric transcription factor CLOCK:BMAL1 (Cox and Takahashi, 2019). In addition to transcriptional regulation, the steady-state levels of rhythmic transcripts are regulated post-transcriptionally (Kojima et al., 2011; Preussner and Heyd, 2016; Wang et al., 2018a). Rhythmic gene expression is critical for the temporal organization of biological functions over the course of the day and night, and its dysregulation by genetic or environmental cues leads to a wide range of pathological disorders including metabolic syndrome, cancer, and cardiovascular disorders (Chaix et al., 2018; Lamia et al., 2008; Rudic et al., 2004; Shimba et al., 2005). While analysis of standard RNA-Seq datasets have been used to investigate if APA regulates cycling transcriptomes (Gendreau et al., 2018; Liu et al., 2013), it still remains unknown if APA isoforms of rhythmically expressed genes are all rhythmically regulated, or if genes characterized as arrhythmically expressed may have one or more cycling APA isoforms.

In this study, we investigated if APA leads to widespread differential expression of distinct transcript isoforms by performing a comprehensive analysis of the diurnal mouse liver transcriptome using 3’ mRNA-Seq. Our results demonstrate that almost a thousand genes in the mouse liver exhibit differential APA transcript rhythmicity, and that hundreds of genes characterized as arrhythmically expressed harbor a rhythmic APA isoform. The underlying mechanisms involve differential post-transcriptional regulation between APA isoforms for about half of these genes, and, to a lesser extent, cell subtype-specific APA isoform expression. Our results also indicate that changes in the transcriptional activity of transcription factors regulate rhythmic gene expression in a largely APA isoform-specific manner, likely through the co-transcriptional recruitment of RBPs that regulate mRNA cleavage and polyadenylation. Interestingly, this regulation occurs primarily on distal APA isoforms, and is the basis for 24-hour rhythm in 3’ UTR length for hundreds of genes. Together, our findings demonstrate that the regulation of rhythmic gene expression is largely APA isoform-specific, and suggest that this regulation may have a large impact on the rhythmic regulation of biological functions by the circadian clock.

## Results

### APA isoforms exhibit differential rhythmicity in mouse liver

To determine if APA leads to isoforms having different expression profiles, we investigated APA usage in mouse liver collected across the 24-hour day using 3’ mRNA-Seq (Figure 1A). As previously reported (Zhang et al., 2014), quantification of mRNA expression across gene models showed that almost 30% of the expressed mouse liver transcriptome is rhythmic (Figure 1A, S1A, S1B). To assess whether some genes exhibit rhythmic or arrhythmic expression of APA isoforms, we generated a database of polyadenylation sites (PAS) in mouse liver using 3’ mRNA-Seq reads from over 100 biologically distinct libraries and comprising more than one billion uniquely mapped reads (see methods). We identified 29,199 high-confidence PAS located in 10,160 genes (Figure S1C; Tables S1, S2). Among these genes, 7,693 (75.7%) had 2 or more PAS (Figure S1D).

**Figure 1:**
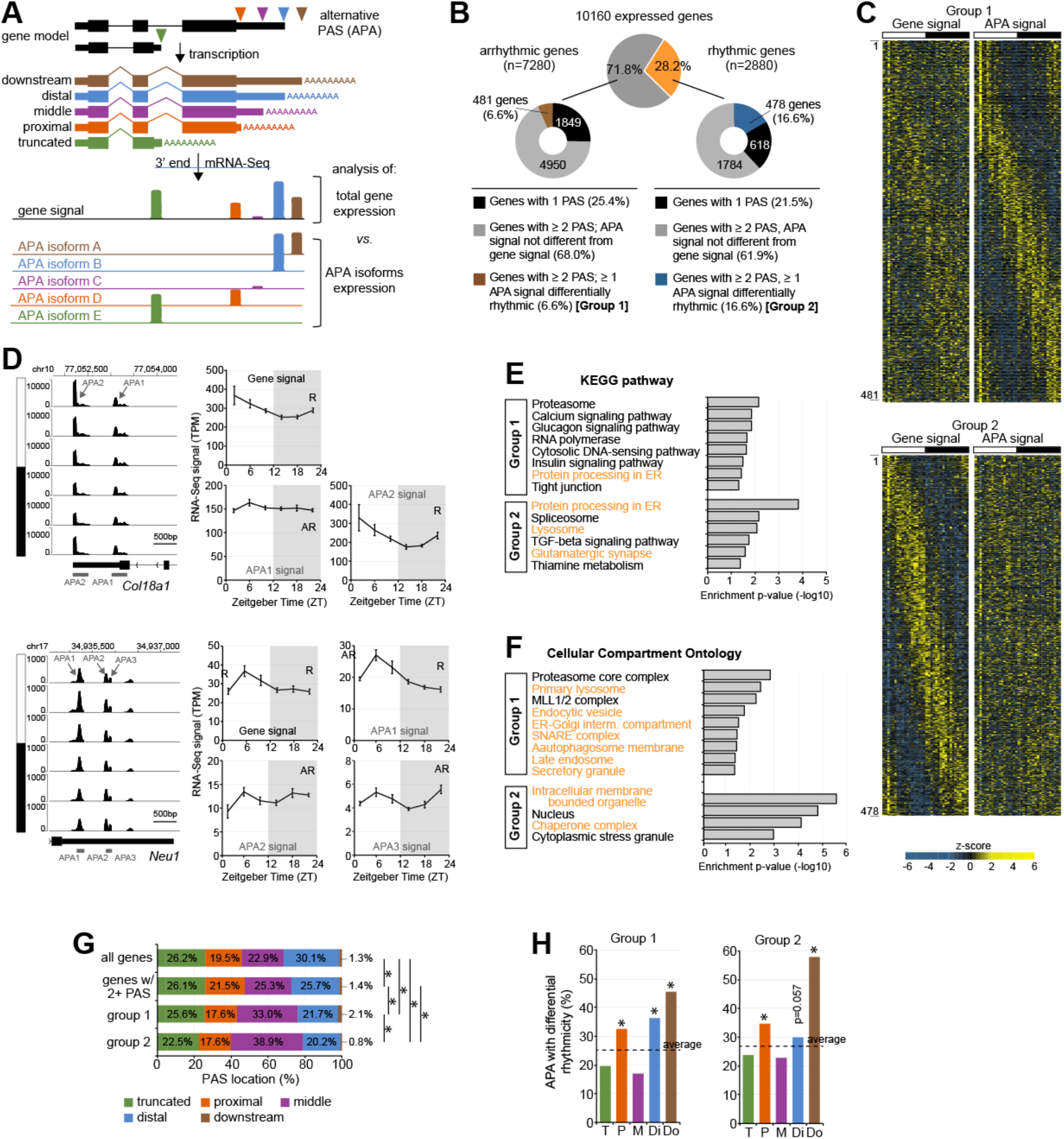
APA isoforms exhibit differential rhythmicity in mouse liver. **(A)** Diagram illustrating how gene transcription can lead to the expression of multiple APA isoforms, and how 3’ mRNA-Seq can be used to analyze the expression of individual APA isoforms. **(B)** Mouse liver 3’ mRNA-Seq rhythmicity analysis, and identification of genes having APA isoforms being differentially rhythmic. Group 1: genes being arrhythmically expressed, but having at least one APA isoform being rhythmic; Group 2: genes being rhythmically expressed, but having at least one APA isoform being arrhythmic. **(C)** Heatmaps representing the expression of a gene (left) and of its corresponding differentially rhythmic APA isoform (right) for Group 1 and 2 genes. Each heatmap column represent a single liver sample (n=36 total; 6 timepoints, n=6 per time point). **(D)** Left: IGV browser view of *Col18a1* and *Neu1* expression across the 24-hour day. Signal for each time point corresponds to the average of 6 biological replicates. Arrows indicate the different APA isoforms, and the day:night cycle is represented by the white and black bars, respectively. Right: expression of *Col18a1* and *Neu1* at the gene and APA isoform levels. R = rhythmic expression; AR = arrhythmic expression. The sum of all PAS signal does not exactly match gene signal because of the normalization procedure (see methods for details). **(E, F)** KEGG pathway (E) and Gene Ontology - Cellular Compartment analysis (F) for genes in Group 1 and 2. **(G)** Distribution of the PAS location for all geens, only those having 2 or more PAS, and for the Group 1 and 2 genes. * denotes significant differences (chi-square test; p < 0.05). **(H)** Enrichment of each PAS location for the differentially rhythmic APA isoforms of Group 1 and 2. * denotes a significant enrichment (hypergeometric test; p < 0.05).

We used this PAS database and combined two statistical analyses to determine if APA isoforms from the same gene can exhibit differences in rhythmic expression. First, we compared rhythmic expression between each gene and their respective APA isoforms. We found that PAS rhythmicity followed for the most part gene rhythmicity, with the majority of arrhythmic genes containing arrhythmic APA isoforms and the majority of rhythmic genes containing rhythmic APA transcripts (Figure S1E). However, PAS rhythmicity did not consistently match the rhythmicity of its corresponding gene, and most rhythmic genes harbored a combination of both rhythmic and arrhythmic PAS. Because this comparative analysis of rhythmicity relies on thresholds and only returns differences in rhythmicity for q-values being just below and above threshold, we performed a second analysis of differential rhythmicity between every APA isoforms and their corresponding gene using DODR (Thaben and Westermark, 2016). Using this more stringent analysis, about 20% of the genes with two or more PAS displayed an APA isoform being differentially rhythmic from gene signal. As expected, no difference was found between PAS and gene rhythmicity for genes with one PAS (Figure S1F). We combined the results of these two analyses to identify two groups of differentially expressed APA isoforms (Figure 1B, S1G, S2A-C; Tables S3-S6). The first group (Group 1) consists of 525 APA isoforms representing 481 genes that are arrhythmic yet have a rhythmic PAS, while the second group (Group 2) consists of 599 PAS representing 478 genes that are rhythmic yet have an arrhythmic PAS. Visualization of the differences between gene and PAS signals with heatmaps (Figure 1C, S1H), along with IGV browser signals for two representative genes (*Col18a1 -Collagen Type XVIII Alpha 1 Chain-*, and *Neu1 - Neuraminidase 1-*; Figure 1D), illustrates that APA isoforms from the same gene can exhibit large differences in rhythmic expression. Based on our stringent analysis, these differences are widespread, and account for about 10% of the genes expressed in mouse liver, and 12.5% of the expressed genes having two or more PAS.

Previous studies have demonstrated that the products of APA isoforms differing in their 3’ UTR length but not in their coding sequence can be located in different cellular membrane compartments, *e*.*g*., endoplasmic reticulum (ER) membrane *vs*. plasma membrane (Berkovits and Mayr, 2015). Interestingly, KEGG pathway and cellular compartment ontology analyses revealed that genes in both group 1 and 2 were enriched for genes associated with intracellular membrane compartments, including lysosome, ER-Golgi, autophagosome, and endosome (Figure 1E, 1F; Table S7).

To get insights into the mechanisms that underlie differential APA isoform expression, we mapped and categorized all PAS as distal 3’ UTR, middle 3’ UTR, and proximal 3’ UTR based on their relative location across the 3’ UTR (Figure 1A). We also included a category termed truncated for PAS located upstream the last exon, as this generates a truncated protein upon translation. Moreover, PAS located up to 1 kb downstream the farthest annotated transcription termination site (TTS) were categorized as downstream (Figure 1A). Mouse liver genes exhibited a relatively equal distribution of proximal, middle, distal, and truncated PAS, with the remaining 3% of PAS being located downstream the annotated TTS (Figure 1G). Genes with differential APA isoform expression (groups 1 and 2) were enriched for PAS located in the middle 3’ UTR, mostly because they have on average a higher number of PAS per gene (Figure S1I; Table S8). Differentially rhythmic APA isoforms were enriched for proximal, distal and downstream PAS, suggesting that longer 3’ UTRs are not the sole factor contributing to differentially rhythmic APA isoform expression (Figure 1H).

### Cell subtype specificity contributes to differential APA isoform expression

Because the liver is a heterogeneous tissue, a mechanism that could explain the differential expression between APA isoforms is cell subtype specificity, with differentially rhythmic isoforms of Group 1 and 2 being expressed in different cell subtypes compared to other isoforms. The liver is composed of Kupffer cells and endothelial cells in addition to the dominant population of hepatocytes. Its anatomical organization into lobules generates a gradient of high oxygen and nutrients (pericentral area) to low oxygen and nutrients (periportal area), that also contributes to hepatocyte subtypes carrying out specialized metabolic functions (Figure 2A) (Braeuning et al., 2006; Rappaport et al., 1954). To determine if differentially expressed APA isoforms are expressed in specific cell subtypes, we examined a public mouse liver single-cell RNA-Seq dataset (Halpern et al., 2017). This dataset was generated using MARS-Seq and thus sequenced 3’ mRNA reads, thereby enabling the analysis of APA isoform expression across the different liver cell subtypes.

**Figure 2:**
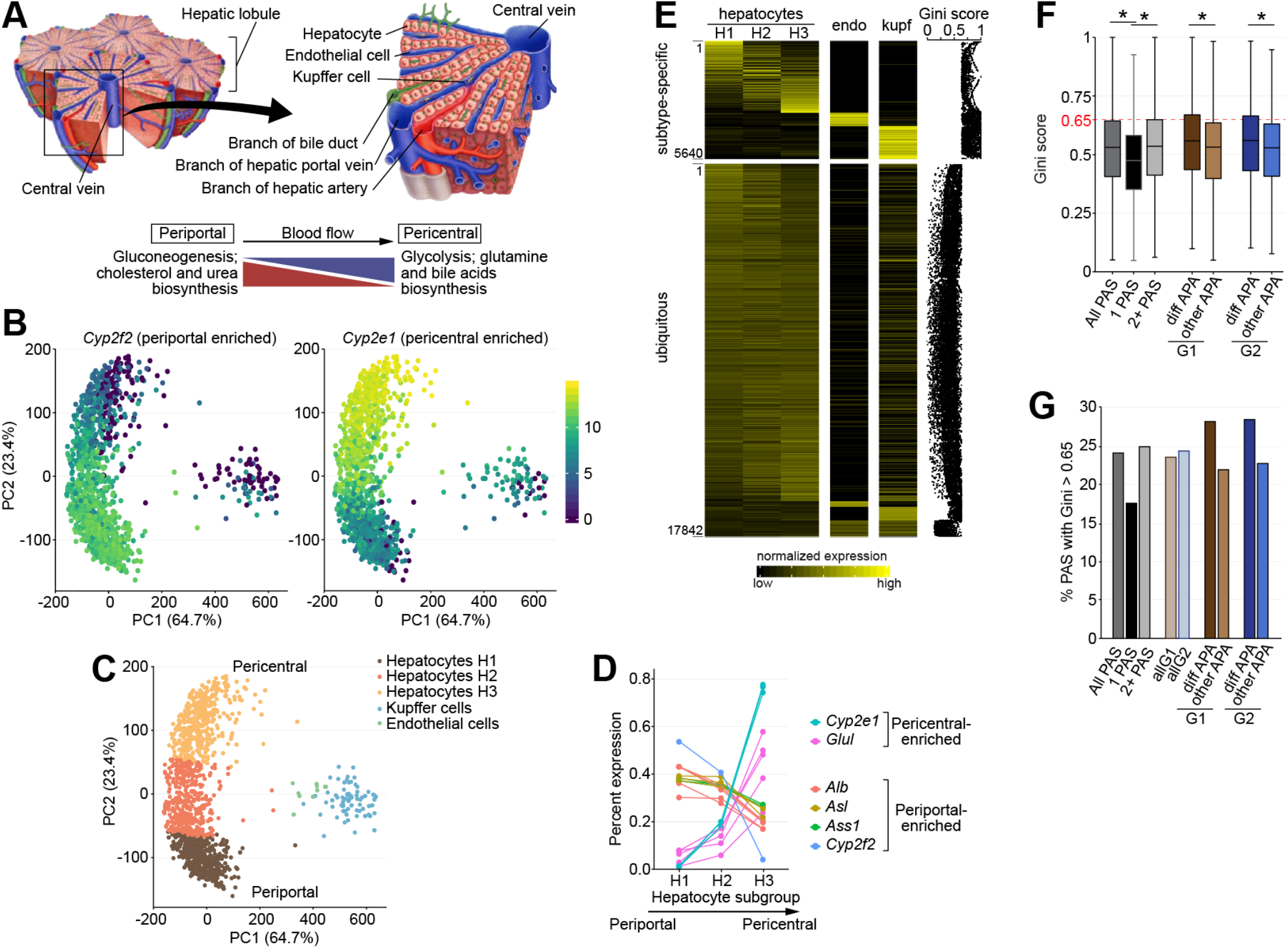
Cell subtype specificity moderately contributes to differential APA isoform expression. **(A)** Anatomical compartmentalization of the liver into lobules and different liver cell types. A gradient from high to low oxygen and nutrient concentration along the periportal-pericentral axis influences the metabolic functions carried out by hepatocytes. **(B)** Principal component analysis (PCA) plot of the distribution of single cells based on the expression of marker genes for hepatocytes, endothelial cells, and Kupffer cells. Cells are shaded according to their expression of *Cyp2f2* (left) and *Cyp2e1* (right). **(C)** Same PCA plot as in B, shaded according to their assigned group. Hepatocyte group H1 is enriched for periportal hepatocytes, while hepatocyte group H3 is enriched for pericentral hepatocytes. **(D)** APA isoform expression of 6 genes used to compartmentalize hepatocytes along a periportal-pericentral axis in the hepatocyte subgroups H1-3. **(E)** Percent expression of each APA isoform in the 3 hepatocyte subgroups, endothelial cells, and Kupffer cells. The Gini coefficient for each isoform is shown on the right. APA isoforms are sorted based on their Gini coefficient and the subgroup in which their expression is the highest. **(F)** Gini coefficient for all PAS, for genes with only one PAS, for genes with 2 or more PAS, and for APA isoforms in Group 1 and 2. diff PAS: differentially rhythmic APA isoforms; other PAS: APA isoforms in Group 1 or Group 2 genes being expressed similarly to the gene signal. * denotes significant differences (Kruskal Wallis test; p < 0.05). **(G)** Percentage of APA isoforms with a Gini coefficient > 0.65.

Several genes preferentially expressed at opposing ends of the liver lobules are commonly used to differentiate hepatocyte subtypes (Braeuning et al., 2006; Gebhardt and Matz-Soja, 2014). Using these hepatocyte marker genes along with marker genes for Kupffer and endothelial cells, we spatially reconstructed the liver subtypes by principal component analysis. Expression analysis of two hepatocyte markers, the cytochrome P450 genes *Cyp2f2* and *Cyp2e1*, across more than 1,000 cells showed biased expression towards the periportal and pericentral ends of liver lobules, respectively (Figure 2B). We used this hepatocyte subtype zonation to split the hepatocyte population into 3 major groups, resulting in a total of 5 groups with the identified Kupffer and endothelial cells (Figure 2C). Subsequent analysis of the APA isoforms for six hepatocyte marker genes confirmed that their expression was partitioned as expected across the liver lobule, thereby validating our spatial reconstruction of the liver using single-cell data (Figure 2D).

To determine if APA isoforms are expressed in specific cell subtypes, we selected the Gini coefficient method as a quantitative measurement of cell subtype specific expression (Kryuchkova-Mostacci and Robinson-Rechavi, 2017; Wright Muelas et al., 2019). This method reports a coefficient ranging from 0 to 1, corresponding respectively to equal expression across all cell subtypes and expression in a unique cell subtype. Unsurprisingly, many APA isoforms only showed small biases in expression across the 5 groups, resulting in Gini coefficients around 0.5 (Figure 2E, S3). Genes with two or more PAS had higher Gini coefficient than genes with one PAS, indicating that APA contributes to cell subtype specific expression. Moreover, differentially rhythmic APA isoforms of Group 1 and 2 exhibited significantly higher Gini coefficients than other transcripts, indicating that subtype specific expression contributes to the differential rhythmic expression of APA isoforms (Figure 2F). To validate this finding, we set a Gini coefficient of 0.65 as a cutoff for cell subtype specific expression, based on the profile of Gini coefficients across all APA isoforms and the Gini coefficient for the different marker genes (Figure S3). Using this cut-off, 28% of the differentially rhythmic APA isoforms of Group 1 and 2 were expressed in a cell subtype specific manner while 22% of the other APA isoforms have a Gini coefficient higher than 0.65 (Figure 2G). This suggests that specific subtype expression has a small, yet significant, effect on the differential rhythmic expression of APA isoforms.

### Post-transcriptional regulation significantly contributes to differential APA isoform rhythmic expression

Given that cell subtype specificity can only partially explain the differential expression of APA transcripts, we next examined if post-transcriptional events could be involved. To this end, we performed 3’ end RNA-Seq of nuclear RNA using the same livers as those used for total 3’ end RNA-Seq. Comparison of the number of intronic reads between total and nuclear RNA, which mostly originate from polyA stretches in introns that are primed by the oligo-dT primer during library first strand synthesis, confirmed that 3’ end RNA-Seq of nuclear RNA increased the number of intronic reads (Figure S4A). Analysis of rhythmic gene expression showed that about a third of the cycling genes do not exhibit rhythmic nuclear RNA expression, indicating as previously reported that post-transcriptional regulation accounts for a fraction of the cycling transcriptome (Figure 3A, 3B; Table S9) (Koike et al., 2012; Menet et al., 2012; Wang et al., 2018a). Additionally, genes found to be rhythmic at both nuclear and total RNA levels, which included all clock genes, displayed similar peak phases of expression (Figure S4B, S4C).

**Figure 3:**
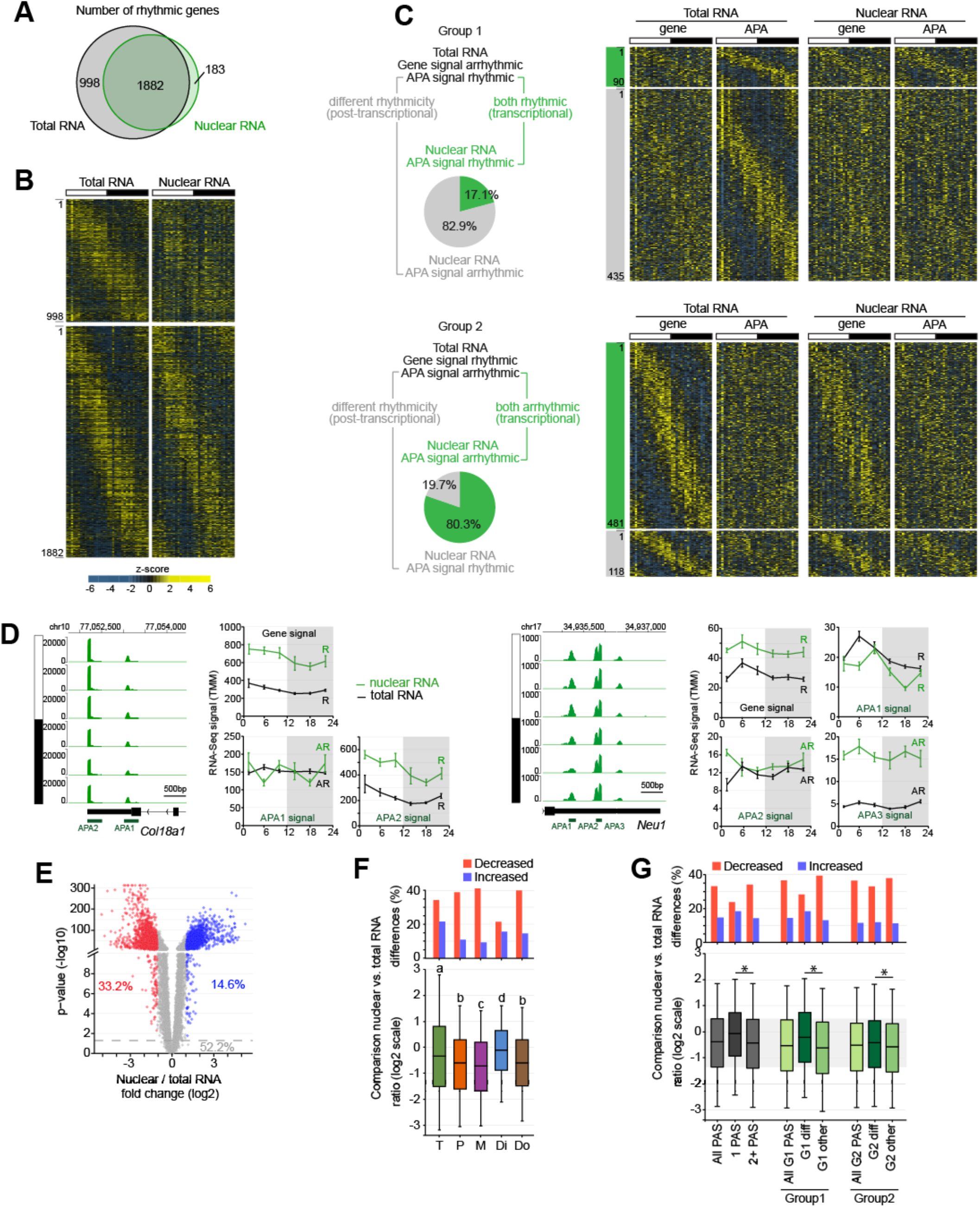
Post-transcriptional regulation significantly contributes to differential APA isoform expression. **(A)** Number of rhythmically expressed genes in total RNA and nuclear RNA. **(B)** Heatmaps representing differential rhythmic gene expression in total RNA (left) and nuclear RNA (right). Each heatmap column represent a single liver sample for both total and nuclear RNA (n=36 total; 6 timepoints, n=6 per time point). **(C)** Pie chart (left) and heatmap (right) representation of the differentially expressed APAS isoforms of Group 1 and 2 in total RNA and nuclear RNA. **(D)** Left: IGV browser view of *Col18a1* and *Neu1* nuclear expression across the 24-hour day. Signal for each time point corresponds to the average signal of 6 biological replicates. Right: expression of *Col18a1* and *Neu1* at the gene level and at the level of individual APA isoforms in total RNA (black) and nucleus (green). R = rhythmic expression; AR = arrhythmic expression. The sum of APA signal does not exactly match gene signal because of normalization procedure (see methods for details). **(E)** Volcano plot representing the log_2_ fold change of nuclear RNA over total RNA against the DESeq2-reported adjusted p-value for all 29,199 PAS (n=36 samples per type of RNA). PAS with a significantly decreased ratio (log_2_ < −1, P_adj_ < 0.05) and significantly increased ratio (log_2_ > 1, P_adj_ < 0.05) are colored in red and blue, respectively. **(F)** Effect of PAS location on the nuclear/total RNA ratio. T: truncated; P: proximal; M: middle; Di: distal; Do: downstream. Groups with different letters are significantly different (Kruskal Wallis test; p < 0.05). **(G)** Distribution of the nuclear/total RNA ratio for APA isoforms of all genes, genes with 1 PAS, genes with 2 or more PAS, and for the differentially rhythmic APA isoforms within G1 and G2. * denotes significant differences (Kruskal Wallis test; p < 0.05).

To determine if post-transcriptional regulation contributes to the differences in APA isoform rhythmicity, we examined the nuclear expression of differentially expressed APA isoforms. Analysis of Group 1 isoforms (rhythmic APA isoforms, arrhythmic genes) revealed that 82.9% of APA isoforms were arrhythmically transcribed (Figure 3C; Table S10), indicating that post-transcriptional regulation largely contributes to their rhythmic expression. Conversely, 80.3% of APA isoforms of Group 2 (arrhythmic APA isoform, rhythmic gene) were transcribed arrhythmically, indicating that 19.7% of these isoforms are post-transcriptionally regulated (Figure 3C). Thus, post-transcriptional events significantly contribute to different APA isoform expression, particularly for those being rhythmically expressed when gene signal is arrhythmic.

Visualization of nuclear signal for *Col18a1* and *Neu1* confirmed these results (Figure 3D). They also revealed large differences in the relative abundance of APA transcripts between nuclear RNA and total RNA with, for example, higher nuclear *vs*. total signal for distal APA isoforms (APA2 for *Col18a1* and APA3 for *Neu1*; Figure 3D). Differences in the relative amount of RNA transcripts in the nucleus *vs*. cytoplasm and/or total RNA are frequently used as a proxy for RNA half-life, with high ratio indicative of short RNA half-life (Luck et al., 2014; Wang et al., 2018a). Differential analysis of nuclear/total RNA ratio with DESeq2 for all APA isoforms revealed widespread differences, with 14.6% PAS exhibiting a high nuclear/total ratio and 33.2% PAS exhibiting a low ratio (Figure 3E). Consistent with our observation with *Col18a1* and *Neu1*, distal APA isoforms had increased nuclear/total signal compared to proximal and middle APA transcripts, suggesting that APA isoforms with longer 3’ UTR have on average shorter half-lives (Figure 3F, Table S11).

To determine if RNA half-life accounts for some of the post-transcriptional regulation of rhythmic APA isoform expression, we examined the nuclear/total RNA ratio of Group 1 and 2 transcripts. Differentially expressed APA isoforms of Group 1, but not of Group 2, displayed increased nuclear/total RNA ratio when compared to other isoforms (Figure 3G). This suggests that shorter RNA half-life likely contributes to the rhythmicity of APA transcripts that are transcribed arrhythmically across the 24-hour day.

### *Bmal1* regulates the expression of many genes in an APA isoform-specific manner

Increasing evidence indicate that RNA processing events, including mRNA decay, mRNA subcellular location, and APA, can be regulated by promoter events (Bregman et al., 2011; Haimovich et al., 2013; Hocine et al., 2010; Moore and Proudfoot, 2009; Trcek et al., 2011; Zid and O’Shea, 2014). The primary underlying mechanism involves the recruitment of RBPs by transcription factors (TFs) to cis-regulatory elements, the subsequent loading of RBPs on the C-terminal domain (CTD) of RNA Polymerase II (Pol II), and RBP deposition onto the nascent transcript during transcription elongation and termination. Because APA can be regulated by RBPs (Di Giammartino et al., 2011; Proudfoot, 2016), we sought to determine if TFs could regulate the expression of specific APA isoforms. We reasoned that if APA is regulated co-transcriptionally by TFs, then, knocking out a TF should affect individual/specific APA isoforms. To test this possibility, we performed 3’ mRNA-Seq across the 24-hour day using liver RNA of clock-deficient *Bmal1*^*-/-*^ mice (*i*.*e*., lacking a circadian activator) fed *ad libitum* (n = 36 mice; n = 6 per timepoint), and quantified APA isoform expression.

As expected, the number of rhythmically expressed APA isoforms was dramatically reduced in clock-deficient *Bmal1*^*-/-*^ mice (Figure 4A). Analysis of differential expression with DESeq2 showed that 28.1% of APA isoforms were significantly affected in *Bmal1*^*-/-*^ mice, with a roughly even distribution between up- and down-regulation (Figure 4B). Consistent with a role of *Bmal1* in driving rhythmic gene expression, affected APA isoforms were enriched for direct *Bmal1*^*-/-*^ target genes and rhythmic APA transcripts (Figure 4C). Interestingly, misregulation of APA isoform in *Bmal1*^*-/-*^ mice was strongly influenced by the phase of rhythmic expression in wild-type mice, with down-regulated APA isoforms having a peak expression biased towards the day:night transition and up-regulated ones showing a bias towards night:day transition (Figure 4D). Given that BMAL1 rhythmic DNA binding peaks during the day (Koike et al., 2012; Rey et al., 2011), these results support the notion that BMAL1 has a direct effect on APA isoforms expression, similar to that shown recently at the gene level (Greenwell et al., 2019; Koronowski et al., 2019; Trott and Menet, 2018).

**Figure 4:**
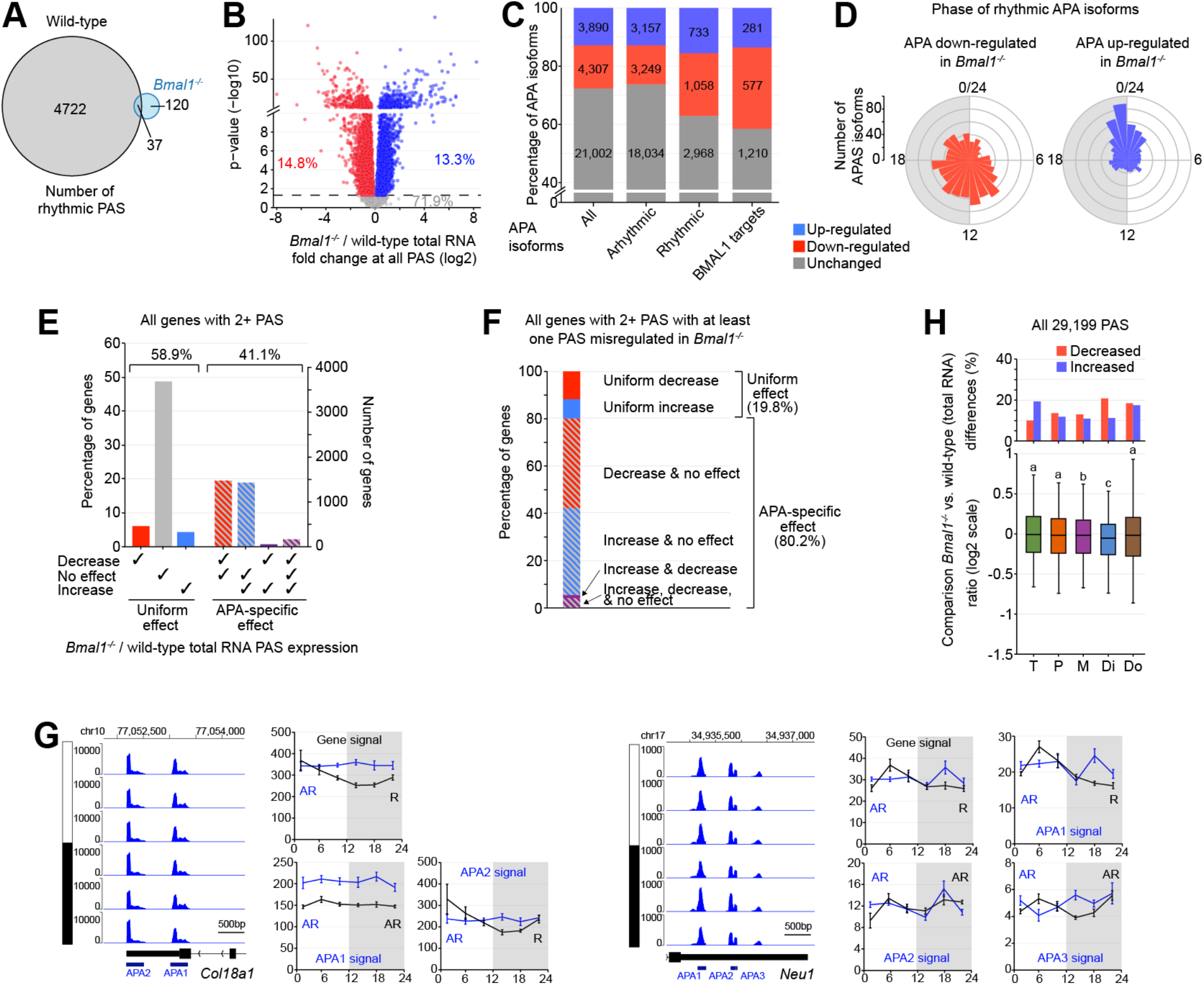
*Bmal1* regulates the expression of many genes in an APA isoform-specific manner. **(A)** Number of rhythmically expressed APA isoforms in wild-type liver total RNA (4722), in *Bmal1*^*-/-*^ liver total RNA (120), or both (37). **(B)** Volcano plot representing the log_2_ fold change of *Bmal1*^*-/-*^/wild-type liver total RNA against the DESeq2-reported adjusted p-value (n=36). **(C)** Number of PAS whose expression is up-regulated, down-regulated, or unchanged between *Bmal1*^*-/-*^ and wild-type mice. **(D)** Peak phase of rhythmically expressed PAS being significantly down- (red) or up-regulated (blue) in *Bmal1*^*-/-*^. **(E)** Number and percentage of genes with 2 or more PAS displaying either a decrease, no effect, an increase, or a combination of effect in *Bmal1*^*-/-*^ mouse liver. **(F)** Percentage of genes with 2 or more PAS and having at least one PAS misregulated in *Bmal1*^*-/-*^ mice displaying either a uniform effect or an APA isoform-specific effect in *Bmal1*^*-/-*^ mouse liver. **(G)** Left: IGV browser view of *Col18a1* and *Neu1* expression in *Bmal1*^*-/-*^ mouse liver across the 24-hour day. Signal for each time point corresponds to the average signal of 6 biological replicates. Right: expression of *Col18a1* and *Neu1* at the gene level and at the level of individual APA isoforms in liver total RNA of wild-type (black) and *Bmal1*^*-/-*^ mice (blue). R = rhythmic expression; AR = arrhythmic expression. The sum of APA signal does not exactly match gene signal because of normalization procedure (see methods for details). **(H)** Effect of PAS location on *Bmal1*^*-/-*^ / wild-type total RNA ratio. T: truncated; P: proximal; M: middle; Di: distal; Do: downstream. Groups with different letters are significantly different (Kruskal Wallis test; p < 0.05).

To determine if APA isoforms from a given gene may be differentially affected by *Bmal1*^*-/-*^, we assayed whether APA isoforms were similarly affected by *Bmal1*^*-/-*^ or if effects were specific to some APA transcripts. We found that 41.1% of the genes exhibited an isoform-specific effect (Figure 4E, S5A). In addition, only 19.8% of the genes having a misregulated APA isoform in *Bmal1*^*-/-*^ exhibited a uniform misregulation among all isoforms (Figure 4F, Table S12). Differences between APA isoform expression in *Bmal1*^*-/-*^ were observed for genes having two or three APA isoforms, indicating that our results were not biased by genes harboring a large number of APA transcripts (Figure S5B). Interestingly, isoform-specific effects in *Bmal1*^*-/-*^ mice consisted almost exclusively of a combination of unaffected APA isoforms with either up-regulated or down-regulated APA transcripts (Figure 4E, 4F). Because an effect of BMAL1 on PAS choice would result in genes having a combination of down-regulated and up-regulated APA isoforms (*e*.*g*., cleavage at proximal PAS instead of distal PAS would concomitantly up-regulate proximal APA isoforms and down-regulate distal APA isoforms), our results strongly suggest that BMAL1 regulates gene expression in an APA isoform-specific manner. This notion is exemplified by *Col18a1*, a BMAL1 direct target gene, in *Bmal1*^*-/-*^ mouse liver (Figure 4G). While the rhythmic distal APA isoform (APA2) exhibits arrhythmic expression in *Bmal1*^*-/-*^ at levels similar to wild-type mice, the proximal APA isoform (APA1) is significantly up-regulated in *Bmal1*^*-/-*^, resulting in an arrhythmic *Col18a1* gene signal in *Bmal1*^*-/-*^ mouse at levels corresponding to the peak expression in wild-type mice (Figure 4G). In contrast, *Bmal1*^*-/-*^ did not affect the expression levels of APA isoforms of *Neu1*, which is another BMAL1 target gene in mouse liver (Figure 4G).

Further investigation revealed that PAS location impacted whether APA isoforms were up- or down-regulated in *Bmal1*^*-/-*^ mice (Figure 4H). Distal APA isoforms showed an overall decreased expression in *Bmal1*^*-/-*^ when compared to wild-type mice (*e*.*g*., lower *Bmal1*^*-/-*^/wild-type expression ratio) whereas truncated and proximal APA isoforms exhibited increased expression in *Bmal1*^*-/-*^ when compared to wild-type mice (as exemplified with *Col18a1*; Figure 4G). Taken together, our results indicate that BMAL1 regulates gene expression in an APA isoform-specific manner, and suggest that BMAL1 promotes the expression of APA isoforms with longer UTRs.

### Rhythmic food intake regulates rhythmic gene expression in an APA isoform-specific manner

To test more specifically if changes in TF activity regulate rhythmic gene expression in an APA isoform-specific manner, we reanalyzed 3’ mRNA-Seq datasets that we generated recently and that investigated how the daily rhythm of food intake, which regulates the activity of metabolic TFs over the course of the day, drives rhythmic gene expression in mouse liver (Greenwell et al., 2019). Specifically, we examined rhythmic APA isoform expression in mice fed either only at night (NR-fed mice) or arrhythmically across the 24-hour day (AR-fed mice). Similar to our findings at the gene level (Greenwell et al., 2019), AR-feeding significantly decreased the number of rhythmically expressed APA isoforms (Figure 5A). Under stringent criteria that included a differential analysis of rhythmicity with DODR, 2,195 APA isoforms were expressed rhythmically in NR-fed mice but not in AR-fed mice (RA group), while only 472 APA isoforms were expressed rhythmically in AR-fed but not in NR-fed mice (AR group) (Figure 5A, 5B). Remarkably, isoforms showing differences in rhythmicity between feeding protocols were often the sole isoform affected for a given gene. Only 18.9% of genes with at least one RA APA isoform had another isoform showing differences in rhythmicity between the feeding protocols (Figure 5C). Similarly, only 25.1% of the genes with at least one AR APA isoform had another APA transcript exhibiting differential rhythmicity (Figure 5C). Importantly, this APA isoform-specific regulation was observed even for genes harboring a large number of APA isoforms, indicating that this observation was not biased by genes with few APA transcripts (Figure 5D, S6A). These results therefore indicate that feeding rhythms affect rhythmic gene expression in a largely APA isoform-specific manner.

**Figure 5:**
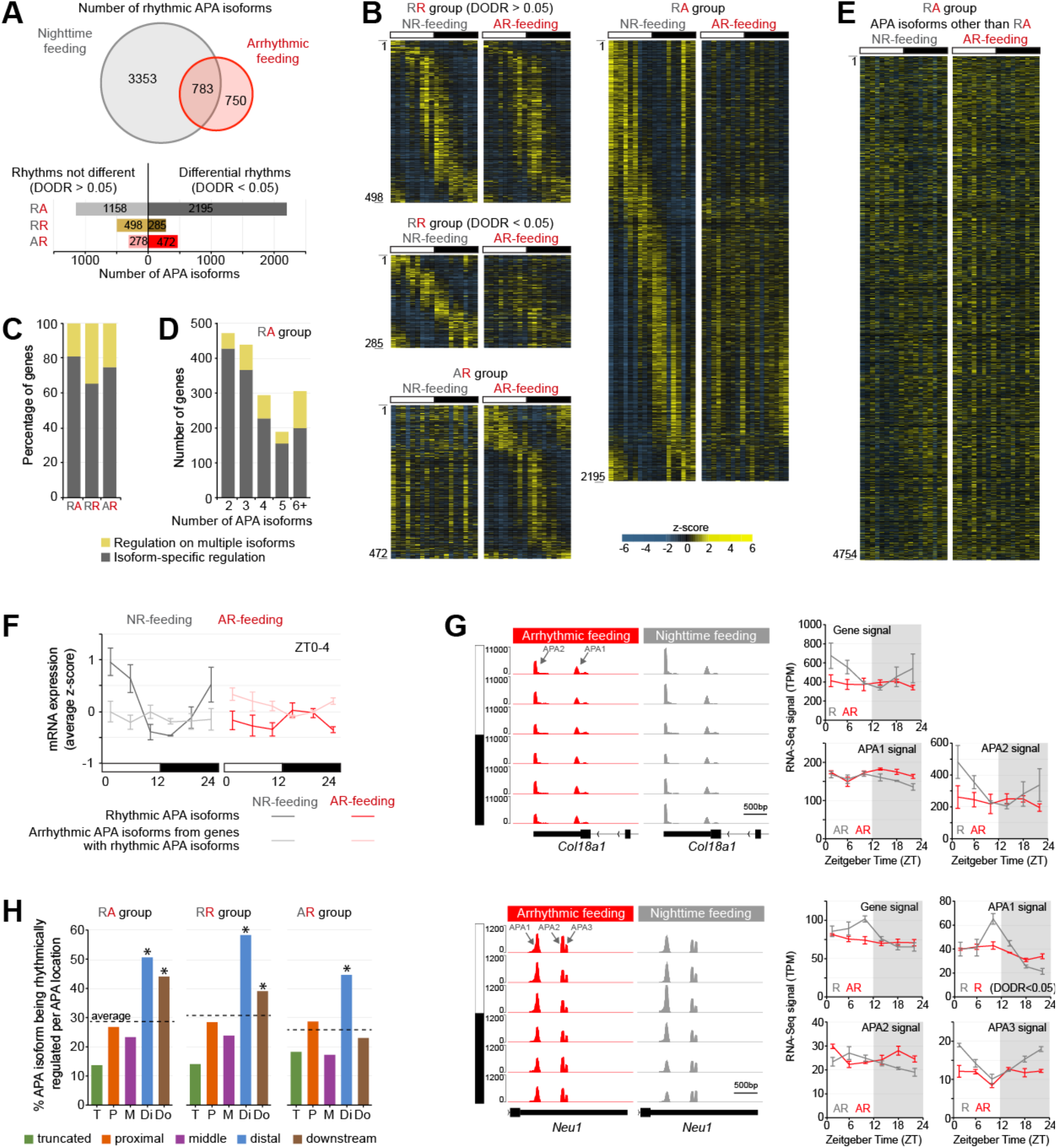
Feeding rhythms regulate rhythmic gene expression in an APA isoform-specific manner. **(A)** Number of APA isoforms rhythmically expressed in both AR-fed and NR-fed mice (RR group), NR-fed mice only (RA group), or AR-fed mice only (AR group). Top: Venn diagram representation. Bottom: number parsed based on differential rhythmicity analysis (DODR, p<0.05). **(B)** Heatmaps representing APA isoforms rhythmic in both AR-fed and NR-fed mice (RR group), NR-fed mice only (RA group), or AR-fed mice only (AR group). Only genes showing differential rhythmicity (DODR, p<0.05) are represented for the RA and AR groups. Each heatmap column represent a single liver sample (n=18 total per condition; 6 timepoints, n=3 per time point). **(C)** Percentage of genes in the RR, RA, and AR groups exhibiting differential rhythmicity for a unique APA isoform (grey) or for multiple isoforms (yellow). **(D)** Number of genes in the RA group exhibiting differential rhythmicity for a single APA isoform or for multiple isoforms, and parsed based on the number of APA isoforms per gene. **(E)** Heatmaps representing the arrhythmically expressed APA isoforms located in a gene containing an APA transcript being rhythmic in NR-fed mice only (RA group). APA isoforms are ordered based on the phase of the rhythmic APA transcript located in the same gene. Only genes showing differential rhythmicity (DODR, p<0.05) were considered. Each heatmap column represent a single liver sample for AR- and NR-fed mice (n=18 total per condition; 6 timepoints, n=3 per time point). **(F)** Expression of APA isoforms being rhythmic in NR-fed mice only (RA group) with a peak expression occurring between ZT0 and ZT4 (dark grey and dark red), and of arrhythmic APA isoforms transcribed from a gene harboring a rhythmic isoform in NR-fed mice only (light grey and pink). Expression corresponds to the averaged z-score +/- sem of three mice per timepoint. **(G)** Left: IGV browser view of *Col18a1* and *Neu1* expression in the liver of NR-fed (grey) and AR-fed (red) mice across the 24-hour day. Signal for each time point corresponds to the average signal of 3 biological replicates. Right: expression of *Col18a1* and *Neu1* at the gene level and at the level of individual APA isoforms. R = rhythmic expression; AR = arrhythmic expression. The sum of APA signal does not exactly match gene signal because of normalization procedure (see methods for details). **(H)** Location of APA isoforms (genes with 2+ PAS) being rhythmic in both AR-fed and NR-fed mice (RR group), NR-fed mice only (RA group), or AR-fed mice only (AR group). * denotes a significant enrichment (hypergeometric test; p < 0.05).

To confirm that our findings were not just due to rhythmicity thresholds and/or stringent analysis, we compared the expression of APA isoforms from the same genes based on their differential expression (*e*.*g*., APA isoforms vs. other APA isoforms within a gene carrying a rhythmic APA isoform in AR- and/or NR-fed mice). Visualization of these other APA isoform expression in NR-fed mice with heatmaps, along with the computation of the average z-score per bin of 4 hours, confirmed that these other APA isoforms were expressed arrhythmically in NR-fed mice (Figure 5E, 5F, S6B, S6C). Examination of *Col18a1* and *Neu1* expression in AR- *vs*. NR-fed mice exemplified these results (Figure 5G). The difference in *Col18a1* rhythmicity at the gene level between NR- and AR-fed mice was solely due to changes in the level of the distal APA2 isoform, as the proximal APA1 isoform remained arrhythmically expressed in both feeding paradigms. Similarly, the rhythmic expression of *Neu1* gene in NR-fed but not AR-fed mice was due to the differential expression of the proximal APA1 isoform between the two feeding protocols (Figure 5G). Interestingly, the distal APA3 isoform of *Neu1* was also rhythmically expressed in NR-fed mice. However, its peak of expression was in antiphase to that of APA1 isoform, suggesting that the decreased cleavage of the proximal APA1 at dusk contributes to the increase APA3 signal at dawn. Such a mechanism is however not prevalent at the genome-wide level, as most genes harbor only one APA isoform being differentially rhythmic between AR- and NR-fed mice (Figure 5C, 5D).

Finally, we examined the PAS location of differentially rhythmic APA isoforms between NR- and AR-fed mice, and found that they were strongly enriched in distal PAS and depleted in truncated PAS (Figure 5H). Together with our finding that BMAL1 facilitates the expression of APA isoforms with longer UTRs, these results suggest that changes in TF activity at promoters and other cis-regulatory regions preferentially regulate the expression of distal APA isoforms.

### Many genes undergo daily rhythms in 3’ UTR length

Next, we sought to determine if differential rhythmicity between APA isoforms could lead to daily rhythms in 3’ UTR length. By computing the relative 3’ UTR length for each gene and examining its rhythmicity across the 24-hour day, we identified 300 genes exhibiting overt rhythms of 3’ UTR length (Figure 6A). Interestingly, a strong phase bias was observed, with 74.0% of transcripts having longer 3’ UTR between ZT20 and ZT08, *i*.*e*., in anticipation of the daytime rest phase in mice (Figure 6B, 6C).

**Figure 6:**
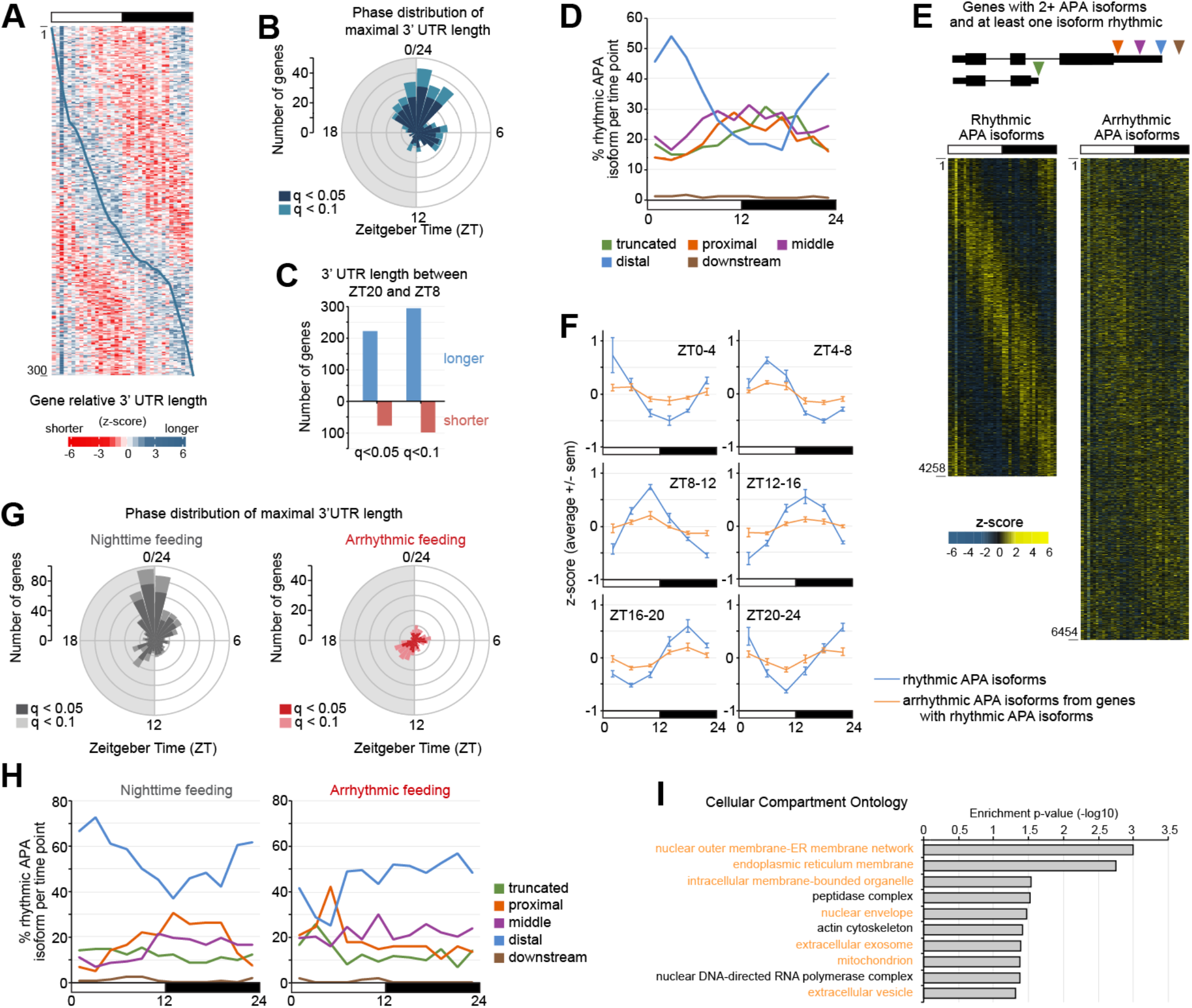
Many genes exhibit a diurnal rhythm of 3’ UTR length. **(A)** Heatmap representing the relative 3’ UTR length for genes showing a significant rhythm in relative 3’ UTR length variation across the 24-hour day (q < 0.05; n = 300). Shorter tail length is displayed in red while longer tail length is displayed in blue. The blue line across the heatmap represents the peak phase of longer relative 3’UTR length. **(B)** Rose plot showing the peak phase of 3’UTR length rhythms (n=300 genes, q<0.05; n=392 genes, q<0.1). **(C)** Number of genes showing a longer (blue) or shorter (red) relative 3’UTR length between ZT20 and ZT08. **(D)** Effect of PAS location on APA isoform peak phase rhythmic expression. Values represent the percentage of rhythmic APA isoform with a specific PAS location exhibiting a peak expression at a specific time of the day per bin of 2 hours. **(E)** Heatmaps representing all APA isoform exhibiting a rhythmic expression in the liver of wild-type mice (left, n=4,258), and representing all arrhythmic APA isoforms located in a gene containing at least one rhythmic APA isoform (right, n=6,454). RNA-Seq signal for each isoform was mean-normalized, and the z-scores calculated for each gene based on all isoforms. Only genes with two or more APA isoforms were included in the heatmaps. For both heatmaps, APA isoforms were ordered based on the phase of the rhythmic APA isoforms. Each heatmap column represent a single liver sample for both total and nuclear RNA (n=36 total; 6 timepoints, n=6 per time point). **(F)** Quantification of APA isoform expression for all rhythmic APA isoforms (blue), and of all arrhythmic APA isoforms located in a gene containing at least one rhythmic APA isoform (orange). Values were calculated as described above for panel E, parsed in bins of 4 hours based on peak expression, and correspond to the averaged z-score +/- sem of 6 mice for each timepoint. **(G)** Rose plot showing the peak phase of 3’UTR length rhythms in NR-fed mice (grey; n=612 genes, q<0.05; n=767 genes, q<0.1) and AR-fed mice (red; n=117 genes, q<0.05; n=186 genes, q<0.1). **(H)** Effect of PAS location on APA isoform peak phase rhythmic expression in NR-fed and AR-fed mice. Values represent the percentage of rhythmic APA isoform with a specific PAS location exhibiting a peak expression at a specific time of the day per bin of 2 hours. **(I)** Gene Ontology - Cellular Compartment analysis for genes showing a rhythm in relative 3’ UTR length with longer UTRs peaking between ZT20 and ZT04 (q < 0.05; n = 166).

To test if the rhythm of 3’ UTR length could be due to isoforms with a specific PAS location peaking at distinct phases, we examined the proportion of distal, proximal, and other APA transcripts that were rhythmically expressed across the 24-hour day per bin of 2 hours. Remarkably, PAS location strongly impacted the phase of rhythmic APA isoform expression, with distal APA transcripts representing the majority of isoforms peaking at dawn, and proximal, middle, and truncated APA isoforms representing a larger fraction of isoforms peaking at dusk (Figure 6D). Because this finding further supports the notion that not all APA isoforms of a gene are rhythmic, we compared rhythmic APA isoforms *vs*. arrhythmic APA transcripts located within genes containing a rhythmic APA isoform using heatmaps and averaged z-score per 4-hour bin. These analyses confirmed striking differences in the rhythmic expression of APA isoforms of a given gene, thus confirming that rhythmic gene expression is largely APA isoform specific (Figure 6E, 6F).

We then tested if feeding rhythms could impact 3’ UTR length by examining the relative 3’ UTR length in NR-fed and AR-fed mice. Similar to the rhythm observed in *ad libitum* fed mice, many genes in NR-fed mice exhibited a rhythm in 3’ UTR length, with longer 3’ UTRs at the end of the night/beginning of the day (Figure 6G). Remarkably, very few genes exhibited a rhythm in 3’ UTR length in AR-fed mice, and no phase preference was observed (Figure 6G). Consistent with the rhythm in 3’ UTR length in NR-fed but not AR-fed mice, distal APA transcripts represented the majority of isoforms peaking at the night:day transition in NR-fed mice, while proximal and middle APA isoforms represented a larger fraction of the APA isoforms peaking at the day:night transition (Figure 6H). Moreover, PAS location had no effect on the phase of rhythmic APA isoform expression in AR-fed mice (Figure 6H). Taken together, these results suggest that rhythmic food intake significantly contributes to the rhythms of 3’ UTR length, and in the location of APA isoform-specific rhythmic expression.

Because alternative 3’ UTRs have been demonstrated to act as scaffolds to regulate membrane protein localization (*e*.*g*., plasma membrane vs. endoplasmic reticulum membrane; Berkovits and Mayr, 2015), we performed a gene ontology and KEGG pathway analysis of the genes displaying a rhythm in relative 3’ UTR length with longer UTRs peaking at dawn (Figure 6I, Table S13). Interestingly, most of them were enriched for proteins associated with membrane-bound organelles, including endoplasmic reticulum, nuclear envelope, and mitochondria (Figure 6I, Table S13). Taken together, these results suggest that APA may generate rhythms of protein expression in specific membrane-bound organelles.

### Rhythmic APA isoforms exhibit a rhythmic enrichment for RBP motifs at their PAS

Increasing evidence indicates that recognition of PAS sequences by the cleavage and polyadenylation (CPA) complex is strongly regulated by *cis*-acting regulatory RBPs that either facilitate or inhibit the usage of proximal PAS (Mayr, 2017; Shi and Manley, 2015). For example, binding of the RBPs HuR (aka *Elavl1*) and TIA1 in 3’ UTRs favors the usage of distal PAS (Hilgers et al., 2012; Zheng et al., 2018), whereas CPEB1 binding shortens UTRs (Bava et al., 2013). To determine if the out-of-phase usage between distal and other PAS (Figure 6D) could be regulated by the differential recruitment of RBPs across the 24-hour day, we performed a motif analysis for RBPs at PAS of rhythmically expressed APA isoforms (−150bp to + 50bp from PAS). To control for background motifs, we used the 6,454 arrhythmic PAS located within genes containing at least one rhythmic isoform (Figure 6E). In addition, we parsed the 4,258 rhythmic PAS into six equally-sized bins based on APA isoform peak expression to eliminate the effects of bin size on p-value enrichment (Figure 7A). Using these parameters for the analysis of 84 unique mammalian RBP motifs, we found that more than half of the RBP motifs were enriched in at least one of the six bins (45 out of 84; Figure 7A, Table S14). Strikingly, almost all RBP motifs showing enrichment in at least one bin displayed an enrichment that is rhythmic over the light:dark cycle (Figure 7B, Table S14). RBP motif enrichment was mostly concentrated towards specific phases, with for example, 21 RBP enriched motifs peaking between ZT0 and ZT4, *i*.*e*., concomitant to the phase of distal APA isoform expression (Figure 7B, 6D).

**Figure 7:**
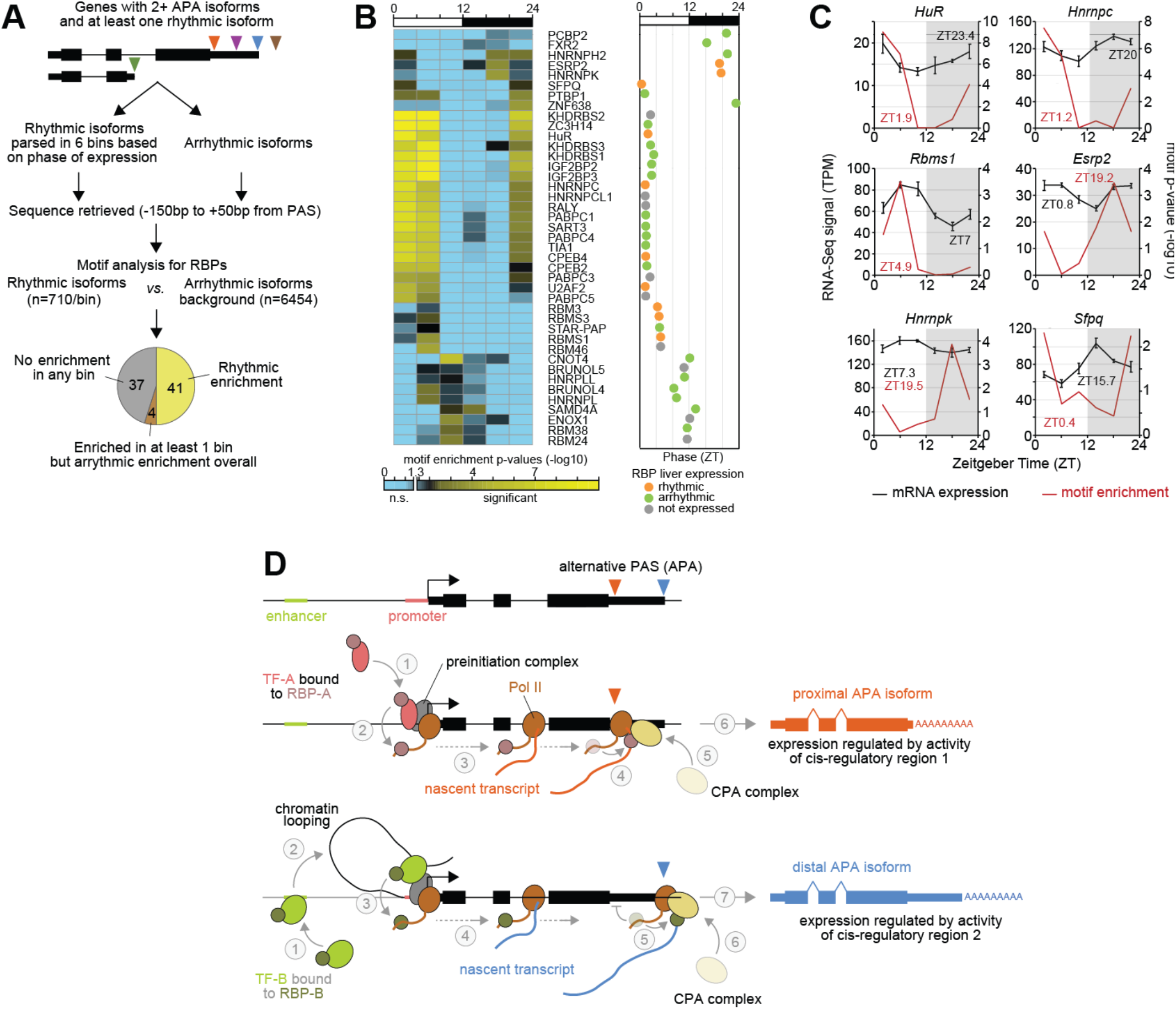
Rhythmic APA isoforms exhibit a rhythmic enrichment for many RBP motifs. **(A)** Flowchart describing the procedure undertaken for the RBP motif analysis at rhythmically expressed APA isoforms, and pie chart illustrating the number of RBPs showing no enrichment at any timepoint (grey; n=37), enrichment in at least one timepoint but with arrhythmic enrichment overall (brown, n=4), and enrichment in at least one time point with rhythmic enrichment (n=41, yellow). **(B)** Left: Heatmap illustration of the rhythmic enrichment for 41 unique RBP binding motif (blue: no enrichment; yellow: high enrichment). Right: Phase of rhythmic enrichment for each of the 41 RBP binding motif. The color of each dot illustrates the RBP mRNA expression in mouse liver: orange: rhythmic expression; green: arrhythmic expression; grey: no expression. **(C)** Mouse liver mRNA expression (black) and RBP binding motif enrichment (maroon) for six rhythmically expressed RBPs displaying a rhythmic motif enrichment. Peak phase of mRNA expression and motif enrichment are displayed in black and maroon, respectively. **(D)** Model describing how transcription factor-driven activity of cis-regulatory regions can regulate the loading of RBPs on nascent transcripts and, consequently, regulate the expression of specific APA isoforms (see text for details). CPA complex: cleavage and polyadenylation complex; TF-A and TF-B: transcription factor A and B, respectively; RBP-A and RBP-B: RNA binding protein A and B, respectively.

Of the 41 RBPs whose binding motifs displayed a rhythmic enrichment, 10 were expressed rhythmically at the mRNA level in mouse liver, while 22 were expressed arrhythmically and 9 were not expressed (Figure 7B). Interestingly, the phase of rhythmic RBP expression coincided for several of them with their rhythmic motif enrichment, suggesting that their rhythmic expression likely contributes to the rhythmic expression of their target isoforms (*e*.*g*., HuR, Figure 7C). Conversely, the expression of the two RBPs *Sfpq* and *Hnrnpk* was in antiphase with their motif enrichment (Figure 7C). This may be explained by reports showing that the binding of these RBPs to 3’ UTRs facilitates miRNA recruitment and downstream degradation of their target mRNA (Bottini et al., 2017; Li et al., 2019). However, the lack of rhythmic expression for most RBPs suggests that rhythmic RBP expression alone does not account for their potential rhythmic binding to 3’ UTRs, and that other mechanisms such as co-transcriptional RBP loading to Pol II may be in play.

Taken together, our analysis demonstrates that the 3’ UTRs of rhythmic APA isoforms are enriched for a large number of RBP motifs, and suggest that the rhythmic binding of RBPs at specific PAS significantly contributes to isoform-specific regulation of rhythmic gene expression in mouse liver.

## Discussion

Rhythmic gene expression is critical for the temporal organization of nearly every biological function across the 24-hour day, and its disruption often leads to the development of pathological disorders (Bass and Takahashi, 2010). Cycling transcriptomes have been characterized in many species and tissues, with RNA-Seq datasets predominantly analyzed as in other fields by concatenating the different transcript variants in a single gene model. Yet, increasing evidence indicate that different isoforms from a single gene can carry out different functions. In the case of APA, which leads to truncated transcripts or transcripts with different 3’ UTR lengths, isoforms can have different stability, subcellular localization, and translation efficiency (Tian and Manley, 2013, 2017). In the present study, we showed that most rhythmically expressed genes contain a combination of rhythmic and arrhythmic APA isoforms, and that hundreds of genes categorized as arrhythmic actually harbor rhythmic APA isoforms. Isoforms peaking at dusk are surprisingly depleted in distal APA transcripts, thereby contributing to the expression of isoforms with shorter 3’ UTRs in anticipation of the mouse active phase. Taken together, our findings indicate that rhythmic gene expression is largely APA isoform-specific, thereby uncovering an integral role of APA in the circadian clock control of cellular functions.

Differences in rhythmic expression between APA isoforms does not appear to rely on a single mechanism, but rather seems to involve several independent processes. While many transcript isoforms are expressed in specific cell subtypes (*e*.*g*., 24.4% in mouse liver; Figure 2) (Booeshaghi et al., 2020; Hu et al., 2020; Lianoglou et al., 2013), cell subtype-specificity only contributes moderately to the differential rhythmicity between of APA isoforms. In contrast, post-transcriptional regulation is highly prevalent, especially for the generation of rhythmic APA isoforms from arrhythmically transcribed transcripts (*i*.*e*., Group 1 isoforms, Figure 3). The contribution of post-transcriptional events to rhythmic gene expression has been widely reported (Koike et al., 2012; Kojima and Green, 2015; Menet et al., 2012; Wang et al., 2018a), and our results therefore indicate that post-transcriptional regulation extends to the expression of individual transcript isoforms.

Besides post-transcriptional events and cell subtype-specific expression, transcription factor-driven expression of specific APA isoforms emerged as a mechanism contributing to the rhythmic expression of specific APA isoforms. Contrary to prevailing models, *Bmal1*^*-/-*^ does not uniformly affect the expression of every isoform of a gene. More than 80% of the genes having a misregulated isoform in *Bmal1*^*-/-*^ mice show differential regulation between APA isoforms (Figure 4E). In addition, changes in feeding rhythms in wild-type mice affect rhythmic gene expression in a largely APA isoform-specific manner, with 75-80% of the affected genes exhibiting an effect on only a single APA isoform (Figure 5C). Although we cannot completely rule out a role for post-transcriptional events in this process, reports from the literature strongly suggest that food intake regulates gene expression by primarily regulating the transcriptional activity of metabolic transcription factors (Desvergne et al., 2006).

How changes in transcription factor activity impact the expression of specific APA isoforms is not well understood. Report that the loading of the RBP ELAV at promoters influences the recognition of alternative PAS suggests that co-transcriptional recruitment of RBPs may be involved (Oktaba et al., 2015). This possibility is further supported by studies demonstrating that promoter events facilitate the co-transcriptional loading of RBPs or effectors to control several RNA processing events, including mRNA decay (Bregman et al., 2011; Shalem et al., 2011; Trcek et al., 2011), mRNA localization (Bertrand et al., 1998; Niedner et al., 2014; Zid and O’Shea, 2014), polyadenylation (Nagaike et al., 2011), mRNA N6-adenosine methylation (Slobodin et al., 2017), and translation (Slobodin et al., 2017; Zid and O’Shea, 2014). Consistent with this possibility, RBPs play a critical role in APA isoform specific expression, by either facilitating or inhibiting the loading of the CPA complex to proximal PAS and, consequently, by either shortening or lengthening the 3’UTR (Gruber and Zavolan, 2019; Mayr, 2017; Shi and Manley, 2015; Xu and Zhang, 2018). Interestingly, many RBPs are recruited to promoters and enhancers in mammalian cells, and this binding often correlates with target gene expression (Van Nostrand et al., 2020; Xiao et al., 2019). Based on these and our data, it is tempting to speculate a model whereby increased activity of a transcription factor mediates the co-transcriptional loading of RBPs to RNA polymerase II and, subsequently, loading of these RBPs to the nascent transcript to either stimulate or inhibit the usage of proximal or middle PAS (Figure 7D). As a result, rhythmic activity of a transcription factor would mediate the rhythmic expression of specific APA isoforms by promoting the rhythmic loading of RBPs to either proximal, middle, or distal PAS (Figure 7D).

The role of RBPs in APA has been widely described (Liu et al., 2013; Shi and Manley, 2015), and it is commonly assumed that their loading to UTRs alters the relative ratio between proximal and distal APA isoform expression without regulating overall transcription levels. As exemplified by *Col18a1* and *Neu1* expression under AR- and NR-feeding (Figure 5G), our data demonstrate that the expression of APA isoforms from a same gene can be regulated independently from one another. Given that defects in APA have been observed in several diseases including cancer (Chang et al., 2017; Masamha and Wagner, 2018), it will be interesting to determine whether alteration of APA isoforms expression in these disease states solely occurs through shifting the usage of distal to proximal PAS, or if increased activity of oncogenic transcription factors also contributes to 3’ UTR shortening (Mayr and Bartel, 2009).

Characterization of APA isoform expression in different cell types and tissues revealed that proliferative cells (*e*.*g*, immortalized cell lines, ES cells, cancer cells, embryonic cells and tissues) tend to exhibit shorter 3’ UTRs whereas differentiated and senescent cells tend to exhibit longer 3’ UTRs (Chen et al., 2018; Lackford et al., 2014; Shepard et al., 2011). Our finding that many genes in mouse liver exhibit a daily rhythm in 3’ UTR length with shorter UTRs in anticipation to the mouse active phase (Figure 6) suggest that this rhythm likely has some physiological relevance for the regulation of hepatic cell biochemistry across the 24-hour day. Interestingly, disappearance of this rhythm in AR-fed mice strongly suggests some coupling between 3’ UTR length and metabolic functions.

Given the emerging role of UTRs in a wide array of processes (Mayr, 2017, 2019), it is highly likely that the rhythmic regulation of APA isoform expression has large impacts on the rhythmicity of biological functions. The recent demonstration that differences in 3’ UTR length can regulate the subcellular localization of many transmembrane proteins (*e*.*g*., endoplasmic reticulum vs. plasma membrane) without altering the protein sequence would suggest that some proteins could be rhythmic in some specific subcellular compartments while arrhythmic in others (Berkovits and Mayr, 2015). This possibility is supported by our findings that differentially expressed APA isoforms and genes exhibiting a rhythm in the relative 3’ UTR length are enriched in membrane-enclosed organelles (Figure 1E, 1F, 6I). 3′ UTRs have also been shown to assist co-translational multiprotein complex assembly, protein oligomerization, and the formation of protein-protein interactions (Chang et al., 2006; Duncan and Mata, 2011; Halbach et al., 2009; Lee and Mayr, 2019). It will be interesting to determine if the increased number of APA isoforms with longer 3’ UTRs at dawn could be involved and facilitate the formation of specific multiprotein complexes. In summary, our data demonstrate that rhythmic gene expression is largely APA isoform-specific, and that the expression of individual APA isoforms can be regulated, at least in the context of biological rhythms, independently from other isoforms. Given that defects in APA are increasingly associated with health risks and diseases (Creemers et al., 2016; Gruber and Zavolan, 2019; Lee et al., 2018; Mayr and Bartel, 2009), our findings will likely be relevant for linking alternative PAS usage to the development of pathological disorders.

## Acknowledgments

We would like to thank Andrew Hillhouse and Texas A&M Institute for Genome Science and Society (TIGSS) for assistance with high-throughput sequencing. We are thankful to Christopher Bradfield for kindly providing the *Bmal1*^*−/−*^ mouse. We also thank Matt Sachs for comments on the manuscript. This work was supported by Texas A&M University startup funds and in part by the National Institutes of Health (R21 AI144454). The laboratory of C.M. is supported by grants from the National Science Foundation (IOS-1754725), the National Institutes of Health (R01 GM124617) and a Klingenstein-Simons Fellowship Award in Neuroscience. The laboratory of D.B-P. is supported by a grant from the National Institutes of Health (R35 GM126966).

## Authors contributions

B.J.G. and J.S.M. conceived and designed the experiments. T.L. and D.B-P contributed to unpublished data that helped frame the overall project. J.R.B. performed all RNA extractions. B.J.G. generated all sequencing libraries, and performed most of the bioinformatics analysis. B.J.G., C.M., and J.S.M. contributed to interpretation of results. B.J.G. and J.S.M. wrote the manuscript, and all authors contributed to editing the final draft.

## Supplementary Tables

Table S1: location of the 29,199 high-confidence PAS used in this study

Table S2: location of the pre-filtered 86,780 PAS in mouse liver

Table S3: localization of the differentially expressed PAS.

Table S4: TMM signal and statistics for all genes

Table S5: TMM signal for all 86,780 PAS

Table S6: statistics for all 86,780 PAS

Table S7: KEGG pathways and Cellular Compartment Ontology analyses of differentially rhythmic APAS isoforms

Table S8: Position of rhythmic PAS (total RNA, wild-type mice)

Table S9: Comparison of rhythmic gene expression between total RNA and nuclear RNA

Table S10: Analysis of post-transcriptional regulation of differentially expressed PAS in Group 1 and 2

Table S11: Nuclear/total RNA ratio and location of APAS isoforms

Table S12: APA isoform-specific regulation of gene expression in *Bmal1*^*-/-*^ mouse liver

Table S13: KEGG pathway, GO-BP and GO-CC analyses of genes exhibiting rhythms in relative 3’ UTR length

Table S14: RBP motif enrichment

## Methods

### Animals

C57BL6/J and *Bmal1*^*-/-*^ mice were raised in-house on a 12 hour light : 12 hour dark cycle (LD12:12), and maintained on *ad libitum* water and food. Mice were anesthetized with isoflurane, decapitated, and the liver collected. The left lateral lobe was cut into three equivalent-sized pieces for RNA processing, with the remainder stored together. All collected tissues were flash-frozen in liquid nitrogen and stored at - 80°C. All animals were used in accordance with the guidelines set forth by the Institutional Animal Care and Use Committee (IACUC) of Texas A&M University (AUP #2019-0222).

### Nuclear RNA isolation

50 to 100 mg of frozen mouse liver tissue was resuspended in 1X PBS and transferred to a 2mL glass homogenizer. Tissues were homogenized with pestle A 6 times and pestle B 4 times. Nuclei were washed twice with the hypotonic buffer, separated by centrifugation at 1,500 g for 2 minutes at 4°C. Nuclei were then resuspended in hypotonic buffer (10 mM Hepes, pH 7.6, 15 mM KCl, 0.15% NP-40, 1 mM DTT, 1 mM PMSF), added on the top of a sucrose cushion (10 mM Hepes, pH 7.6, 15 mM KCl, 0.15% NP-40, 24% sucrose, 1 mM DTT, 1 mM PMSF), and centrifuged at 20,000 g for 10 minutes at 4°C. The supernatant was carefully removed, and the nuclei washed twice with resuspension buffer (10 mM Tris pH7.5, 150 mM NaCl, 2 mM EDTA, 1 mM PMSF), and used immediately for RNA extraction.

### RNA extraction and processing

Total and nuclear RNA were extracted with TRIzol reagent following manufacturer recommendation. Total RNA was extracted from a frozen liver sample (left lateral lobe), while nuclear RNA was extracted from a nuclei pellet. Frozen tissue or nuclei pellet were mixed with 300µL of TRIzol reagent, homogenized with a pellet mixer, and the volume brought to 1mL with 700mL of TRIzol reagent. 500 µl of TrIzol reagent was added to each 500 µl gradient fractions. After TRIzol reagent RNA extraction, RNA was further purified with an acid phenol/chloroform extraction and ethanol precipitation. Each sample was quantified by NanoDrop-1000 and Promega QuantiFluor ssRNA system, and integrity assessed by gel electrophoresis.

### Library generation and sequencing

RNA-Seq libraries were generated using the Lexogen QuantSeq 3’ mRNA-Seq Library Prep Kit following manufacturer instructions, beginning with 2µg of total RNA as starting material. cDNA was PCR-amplified for 12 cycles following manufacturer recommendations for mouse liver tissue. Libraries were multiplexed in equimolar concentrations and sequenced across multiple runs using an Illumina NextSeq 500 (Molecular Genomics Workspace, Texas A&M University, USA).

### Data processing

Sequenced reads were pre-processed with the R package ShortRead (Morgan et al., 2009) to remove the first 12nt, remove low quality bases at the 30 end, trim poly-A tails and embedded poly-A sequences, and remove all reads under 36nt in length. Reads were aligned to the mm10 transcriptome, assembly GRCm38.p4, with the STAR aligner (Dobin et al., 2013) version 2.5.2b with options: `--outSAMstrandField intronMotif–quantMode GeneCounts –outFilterIntronMotifs RemoveNoncanonical`

Secondary alignments were removed with samtools view -F 0×100. Read counts were summarized with the function summarizeOverlaps from the R package GenomicRanges (Lawrence et al., 2013) using options: `mode=IntersectionStrict inter.feature=FALSE`. Libraries were filtered and normalized by library size using the Trimmed Mean of M-values (TMM) normalization (Robinson and Oshlack, 2010) within edgeR (Robinson et al., 2010) using default settings.

### PAS mapping

Initial PAS definition was performed through the combination of two separate analyses. First, all total RNA reads as well as the reads from the 72 samples from (Greenwell et al., 2019) (total of 108 separate libraries) were trimmed to their 3’ most mapped nucleotide, taking into account the CIGAR string. In the first analysis, the 3’ nucleotides were immediately put through peak calling by HOMER (Heinz et al., 2010) to find a broad range of potential APAS using the following options:

~~~
makeTagDirectory -precision 3 -totalReads all –fragLength 1 –keepAll
findPeaks -strand separate -tbp 0 –fragLength
     1 –size 10 -minDist 25 –ntagThreshold
     2 –region
~~~

resulting in 76,018 prospective PAS.

In the second analysis, we attempted to define all APAS that represented the exact end of transcripts. Therefore, all 3’ most nucleotide reads were filtered to only those containing at least 6 consecutive adenine residues at the 3’ end in the original unmodified read, indicating that these reads are directly against the poly(A) tail. Next, we scanned the genome 20nt downstream of the mapped 3’ end of these reads and removed those with 12 or more adenine residues in those 20nt, indicating that they existed due to internal priming events. Reads in both steps were filtered out using custom Perl scripts. Finally, PAS were defined through HOMER using the following options:

~~~
makeTagDirectory -precision 3 -totalReads all
-fragLength 1 –keepAll
findPeaks -strand separate -tbp 0 -fragLength
1 -size 10 -minDist 25 -ntagThreshold 5 –
region
~~~

resulting in 31,837 prospective APAS.

The two sets of APAS were then concatenated and combined to yield a single list using the reduce function from GenomicRanges (Lawrence et al., 2013). Finally, any APAS found to overlap with the mm10 blacklisted regions generated by ENCODE were removed (Amemiya et al., 2019; Yue et al., 2014). A total of 86,780 possible APAS resulted from these steps (Table S2).

### PAS filtering

PAS were annotated to genes using their overlaps with each gene part with summarizeOverlaps from GenomicRanges (Lawrence et al., 2013), with a final annotation performed using a priority list. TSS were defined as the region +0 to +100nt from the annotated TSS. TTS were defined as the region −20nt to +20nt from the annotated TTS. Finally, downstream regions were defined as up to 2kb from the end of the TTS above. Next, we removed all PAS that contained genomic poly(A) stretches that escaped detection in the previous steps. The region from −15 to +5 of the 3’ end of every PAS was analyzed, and those with 12 or more adenine residues were removed. Since the downstream annotation can potentially result in ascribing APAS of a downstream gene to the upstream and unrelated gene, we removed all PAS that were annotated as downstream yet interior of another gene. Raw count values were then normalized by library size using the Trimmed Mean of M-values (TMM) normalization (Robinson and Oshlack, 2010) within edgeR (Robinson et al., 2010) using default settings.

We next examined whether all APAS for each gene accounted for the gene expression seen in total RNA on a log2-scale. Any genes where the sum of PAS expression overlapping the CDS was less than half of its total expression or more than the total expression + 0.5 were removed (Figure S1D). Finally, we examined the expression of each PAS and included in the final filtered list if their contribution was more than 10% of the total gene signal, or if it had a mean TMM value over 5. Any PAS that had a maximum contribution to any transcript under 0.5% of total as well as a mean TMM value under 0.5 was automatically discarded. All leftover PAS between these two ranges were tested for overlap with a publicly available database of APAS (Wang et al., 2018b). APAS from PolyA_DB were extended upstream to a total of 75nt, and any of our APAS that overlapped with those from PolyA_DB were kept. After filtering, a total of 29,199 high-confidence APAS remained (Table S1).

### Rhythmicity and differential rhythmicity analysis

Rhythmicity analysis for every gene and PAS in the 4 paradigms was performed with four algorithms from three programs: F24 (Hutchison et al., 2015; Wijnen et al., 2005), JTK_CYCLE and LS from MetaCycle (Wu et al., 2016), and HarmonicRegression (Hughes et al., 2009). The resulting p-values from all 4 algorithms were combined using Fisher’s method into one p-value, all of which were then adjusted using the Benjamini-Hochberg method (Benjamini and Hochberg, 1995) within the p.adjust function available in base R to control for the false-discovery rate (FDR). Genes with a q-value under 0.05 were considered rhythmic for that paradigm.

Differential rhythmicity analysis was performed with the robustDODR algorithm within DODR (Thaben and Westermark, 2016). Genes and PAS with a p-value less than were considered as differentially rhythmic.

The similarity analysis was performed by ranking the Pearson correlation of two paired sets of expression values in log2 *vs*. the Pearson correlations from 10,000 trials where one set of values was randomized against the other. The paired set of expression values were considered as similarly expressed if the experimental Pearson correlation coefficient was within the top 5% of the 10,000 coefficients calculated through permutations.

### Differential expression analysis

All differential expression analysis was performed using DESeq2 (Love et al., 2014) using default settings.

### Analysis of BMAL1 target genes

Mouse liver BMAL1 target genes were defined as genes containing a BMAL1 ChIP-Seq peak in their locus (from −10kb from the transcription start sites to +1kb to the transcription termination sites), and exhibiting a differential rhythmicity by DODR (p < 0.05) in *Bmal1*^*-/-*^ mice fed only at night when compared to wild-type mice fed only at night. Specifically, we used the list of BMAL1 ChIP-Seq peaks published by Beytebiere et al., 2019 (Table S1; Beytebiere et al., 2019), and the mouse liver RNA-Seq datasets from *Bmal1*^*-/-*^ and wild-type mice fed only night published by Atger et al., 2015 (Atger et al., 2015), and analyzed as in Greenwell et al., 2019 (Table S2; Greenwell et al., 2019).

### Relative 3’UTR length

The distance between the center of a PAS and the 3’ end of the corresponding gene was calculated for all PAS. These distances were then weighted by the percentage signal each PAS contributed to the total gene signal at each timepoint. Rhythmicity calculations were performed on the weighted relative 3’ UTR length as described above for the analysis of rhythmic gene expression.

### RBP motif analysis

Enrichment of RBPs were calculated by taking the position probability matrices (PPMs) provided by the Hughes lab (Ray et al., 2013) and converting them to positional weight matrices (PWMs). Using the R package PWMEnrich (Stojnic and Diez, 2014), background enrichment for each PWM was calculated using the arrhythmic PAS found in all genes that contain both arrhythmic and rhythmic PAS (n = 6,454 PAS). The final enrichment of each PWM was then calculated against the background PAS for all rhythmic PAS (n = 4,258 PAS), and for PAS parsed in 6 equally-sized bins based on peak phase of expression. Bins were equally sized to eliminate the effects of bin size on p-value enrichment. Starting from the light on signal (ZT0), the length of each bin was 3.42, 5.21, 3.87, 3.8, 4.5, and 3.2 hours. Rhythmic analysis of RBP motif enrichment was performed using Metacyle (Wu et al., 2016).

### Single cell mouse liver reconstruction

Methods indicated in the original paper (Halpern et al., 2017) were followed as closely as possible. Single-cell data were downloaded from GEO (GSE84498) and aligned to the mm10 transcriptome as above. Gene expression for all genes as well as the ERCC92 spike-in was performed using summarizeOverlaps from the GenomicRanges package (Lawrence et al., 2013) with the following options:

~~~
mode = “IntersectionStrict”, singleEnd = TRUE,
     ignore.strand = FALSE, inter.feature =
     FALSE
~~~

Cells were removed if the total of their reads mapping to ERCC92 was greater than 4% of the total of reads mapping to the genome (Risso et al., 2014). Genes were removed from consideration if the total number of reads mapping to them was 0. All libraries were then normalized to library depth using the TMM normalization as above. Next, we summed the expression of markers for hepatocytes (*Apoa1, Apob, Pck1, G6pc, Ttr)*, endothelial cells (*Kdr, Egf17, Igfbp7, Aqp1*), and Kupffer cells (*Clec4f, Csf1r, C1qc, C1qa, C1qb*). Cells with a greater total for the endothelial or Kupffer marker genes were labeled as endothelial or Kupffer cells, respectively. Cells that had a higher total for both over the hepatocyte marker genes were discarded. All remaining cells were labeled as hepatocytes. Any hepatocyte cell with less than 1% of total expression coming from albumin (*Alb*) was discarded. In order to separate hepatocytes by their zonation profile, PCA analysis was performed on all hepatocytes using their expression for the 20 marker genes indicated in (Braeuning et al., 2006), and hepatocytes were split into three even groups (H1, H2, H3) based on their PC1 value. Finally, another PCA analysis was performed on all cells using a combination of the Braeuning hepatocyte, Kupffer, and endothelial marker genes.

For PAS expression in the single-cell data, the 3’ end of all reads was taken and quantified across all accepted 29,199 PAS. The raw reads for all 5 groups (H1, H2, H3, Kupffer, and endothelial) were summed up. Due to the differing chemistries between 3’ QuantSeq and MARS-Seq, some PAS did not match well, and so any PAS with fewer than 5 reads in total was removed. The remaining PAS were then TMM-normalized as above.

### KEGG and GO analyses

All KEGG and GO analyses were performed using the kegga and goana functions available within limma (Ritchie et al., 2015). Gene symbols were converted to Entrez IDs with AnnotationDbi (Pagès et al., 2019).

### Access to sequencing datasets

All RNA-Seq datasets have been deposited to Gene Expression Omnibus (GEO). Access is currently password-protected, but datasets will become publicly available under the Series GSE151173 once this manuscript is accepted for peer-reviewed publication.

**Figure S1:**
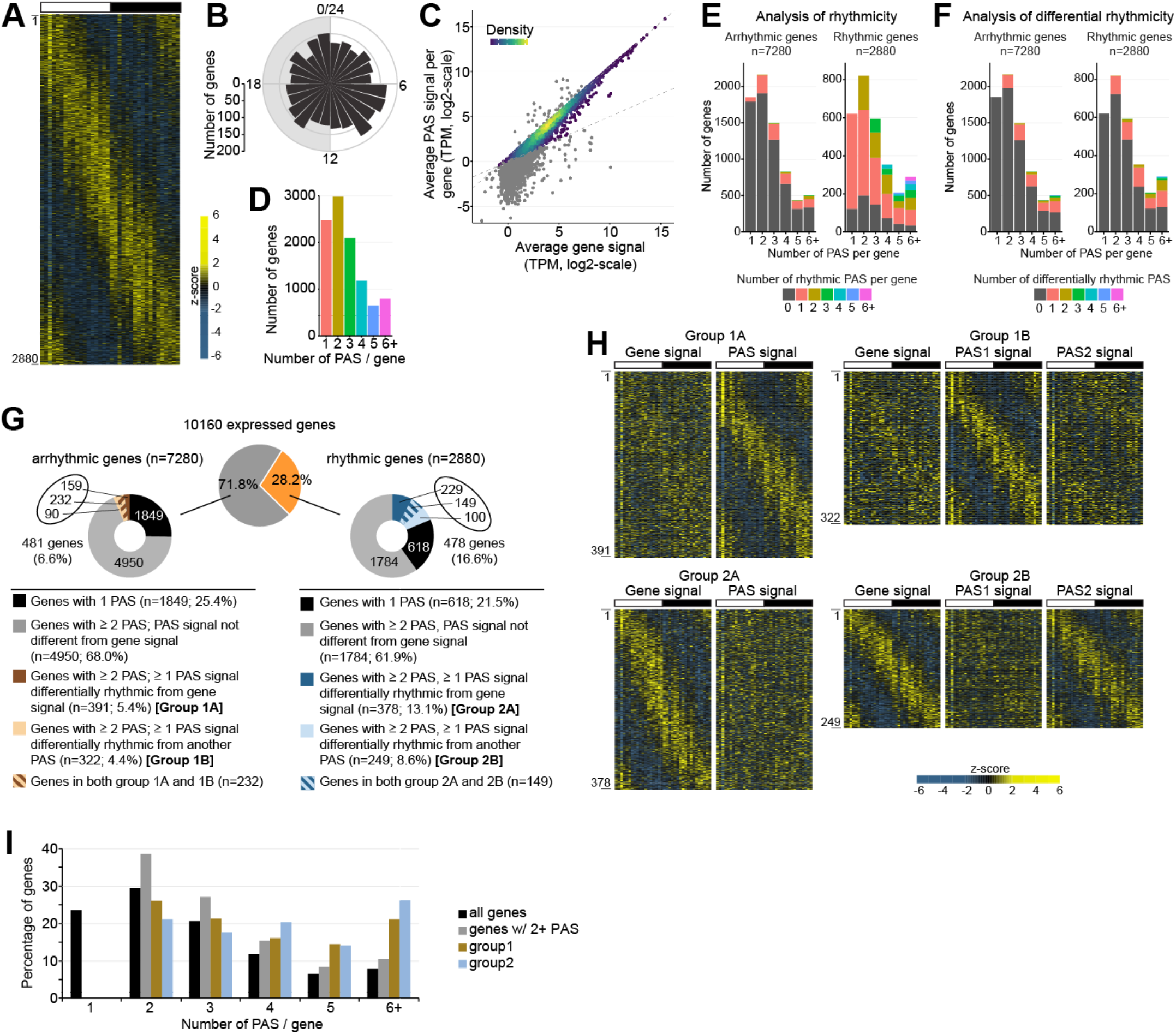
Characterization of rhythmic APA isoform expression in mouse liver (related to Fig. 1) **(A)** Heatmap of the 2,880 genes being rhythmically expressed in mouse liver. Each heatmap column represent a single liver sample (n=36 total; 6 timepoints, n=6 per time point). **(B)** Rose plot representing the distribution of peak phases of rhythmic gene expression. **(C)** Correlation between the expression of a gene and the sum of expression from its APA isoforms. APA isoforms that were excluded (see Methods section for details) are shaded gray. Those colored in purple/green/yellow were used to generate the final list of 29,199 APA isoforms that are considered in the present study. **(D)** Number of PAS per gene (n = 10,160 expressed genes) in mouse liver. **(E)** Rhythmicity of APA isoforms per gene, binned based on the number of APA isoforms per gene. **(F)** Number of differentially rhythmic APA isoforms (based on DODR analysis) per gene, binned based on the number of APA isoforms per gene. **(G)** Rhythmicity analysis used for the identification of genes having differentially rhythmic APA isoforms. This analysis identified 4 groups of genes (1A, 1B, 2A, 2B) that were combined into the two groups G1 and G2 considered in the current manuscript. **(H)** Heatmaps representation of total RNA expression of a gene (left) and of its corresponding differentially rhythmic APA isoform (right) for genes belonging to the groups 1A, 1B, 2A, 2B. **(I)** Number of PAS per gene for all genes, only those having 2 or more PAS, and for the Group 1 and 2 genes.

**Figure S2:**
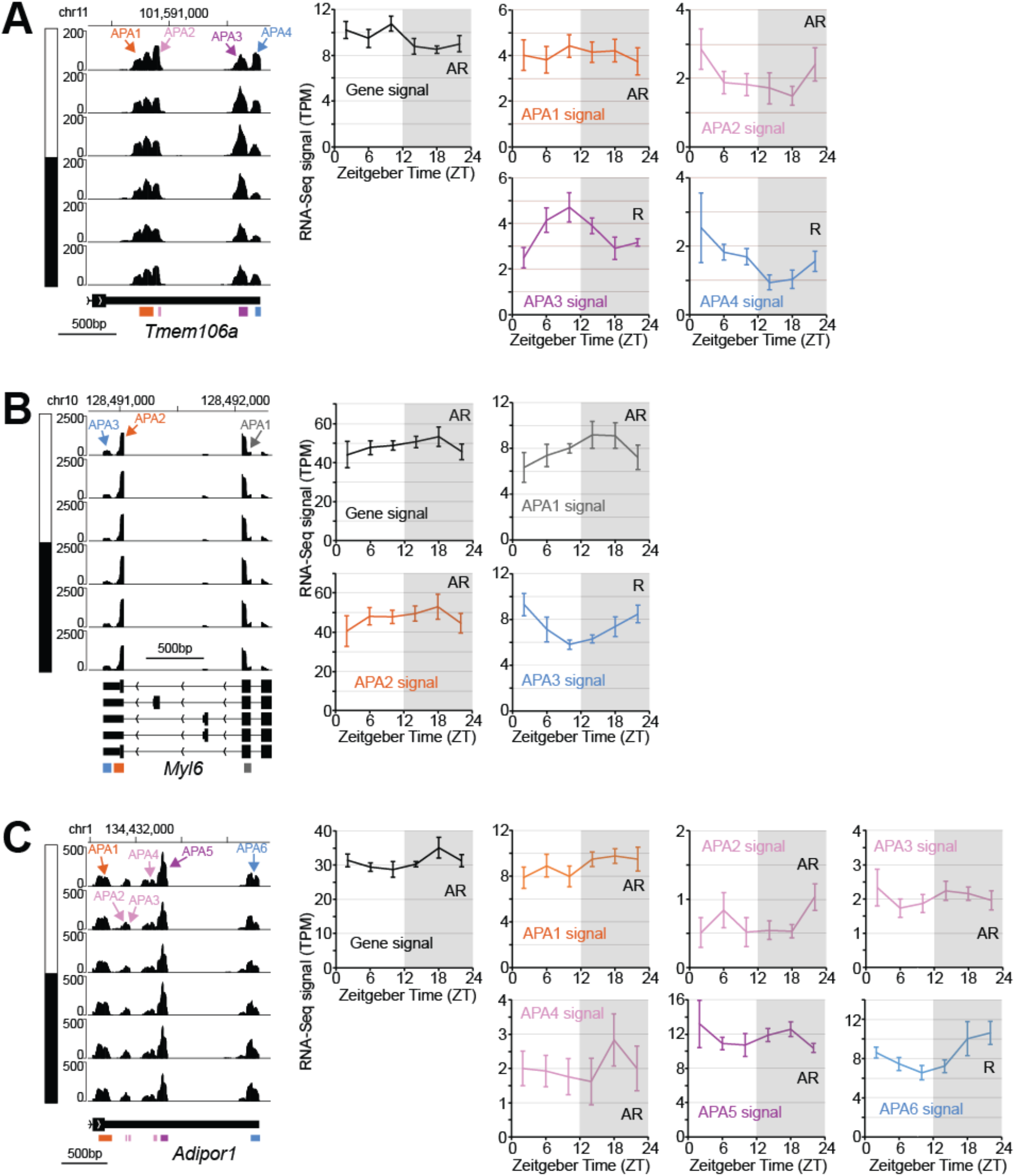
Examples of Group 1 genes (related to Fig. 1) **(A-C)** APA isoform expression for three Group 1 genes, *i*.*e*., genes that are arrhythmically expressed but harbor an APA isoform that is rhythmically expressed. Left: IGV browser view of *Tmem106a* (A), *Myl6* (B) and *Adipor1* (C) expression across the 24-hour day. Signal for each time point corresponds to the average of 6 biological replicates. Arrows indicate the different APA isoforms, and the day:night cycle is represented by the white and black bars, respectively. Right: expression of *Tmem106a* (A), *Myl6* (B) and *Adipor1* (C) at the gene and APA isoform levels. R = rhythmic expression; AR = arrhythmic expression. The sum of all PAS signal does not exactly match gene signal because of the normalization procedure (see methods for details).

**Figure S3:**
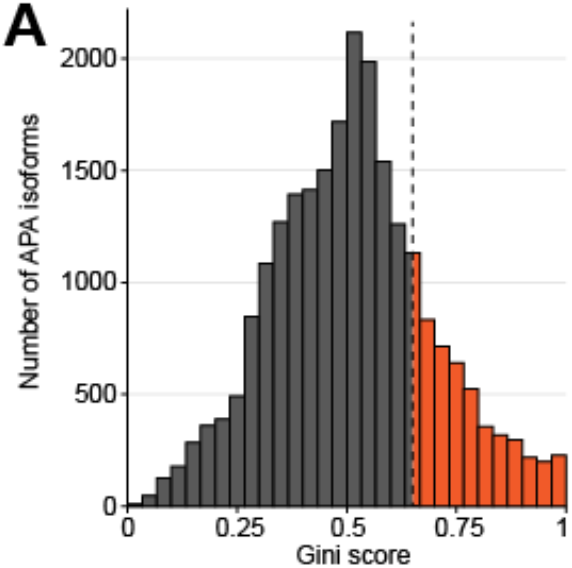
Analysis of mouse liver APA isoform expression in different mouse liver cell subtype. (related to Fig. 2) Distribution of Gini scores for all APA isoforms in mouse liver. Gini scores above 0.65, which were considered as indicators of cell subtype specific expression, are colored in orange.

**Figure S4:**
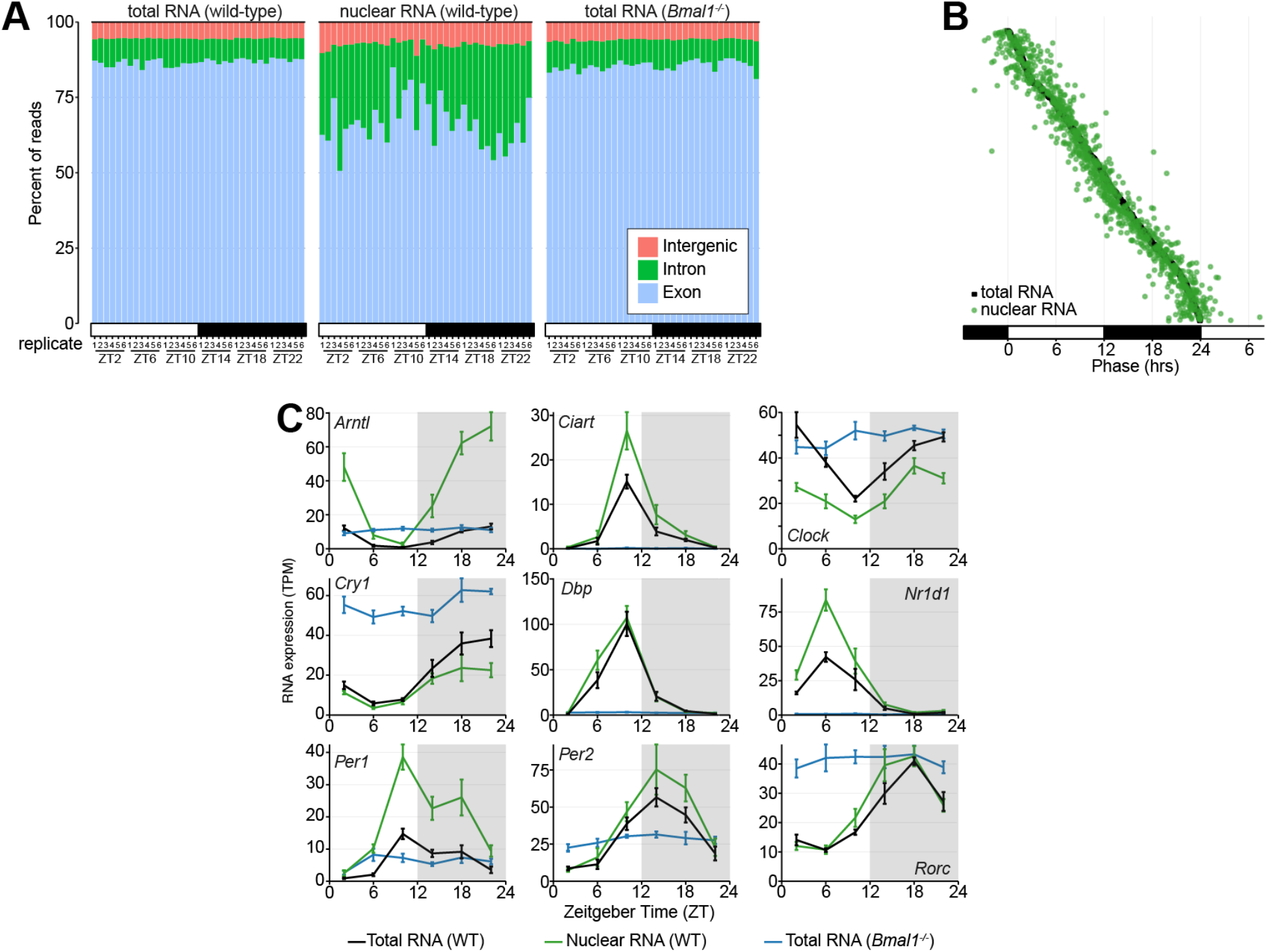
Analysis of mouse liver APA isoform expression in nuclear RNA. (related to Fig. 3) **(A)** Percent of reads aligning to exons, introns, or intergenic regions for the three types of libraries sequenced in this study (n = 108 libraries total). **(B)** Peak phase of rhythmic gene expression in total RNA (black) and nuclear RNA (green) for the genes being rhythmically expressed in both total and nuclear RNA. **(C)** Expression of core clock genes in the liver across the 24-hour day. Black: total RNA, wild-type mice. Green: nuclear RNA, wild-type mice. Blue: total RNA, *Bmal1*^*-/-*^ mice. Each value represents the average +/- sem of 6 mice.

**Figure S5:**
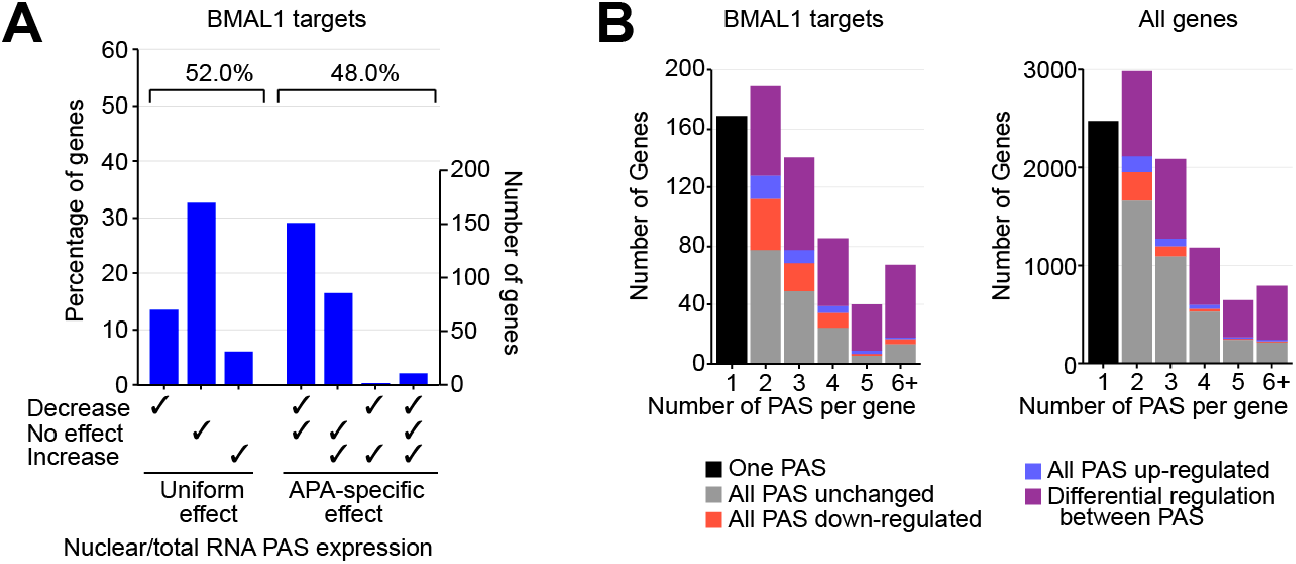
Analysis of mouse liver APA isoform expression in *Bmal1*^*-/-*^ mice. (related to Fig. 4) **(A)** Percentage and number of BMAL1 target genes with 2 or more PAS displaying either a decrease, no effect, an increase, or a combination of effect in *Bmal1*^*-/-*^ mouse liver. **(B)** Number of genes with at least 2 APA isoforms showing either a decrease, no effect, an increase, or a combination of effect *Bmal1*^*-/-*^ mouse liver.

**Figure S6:**
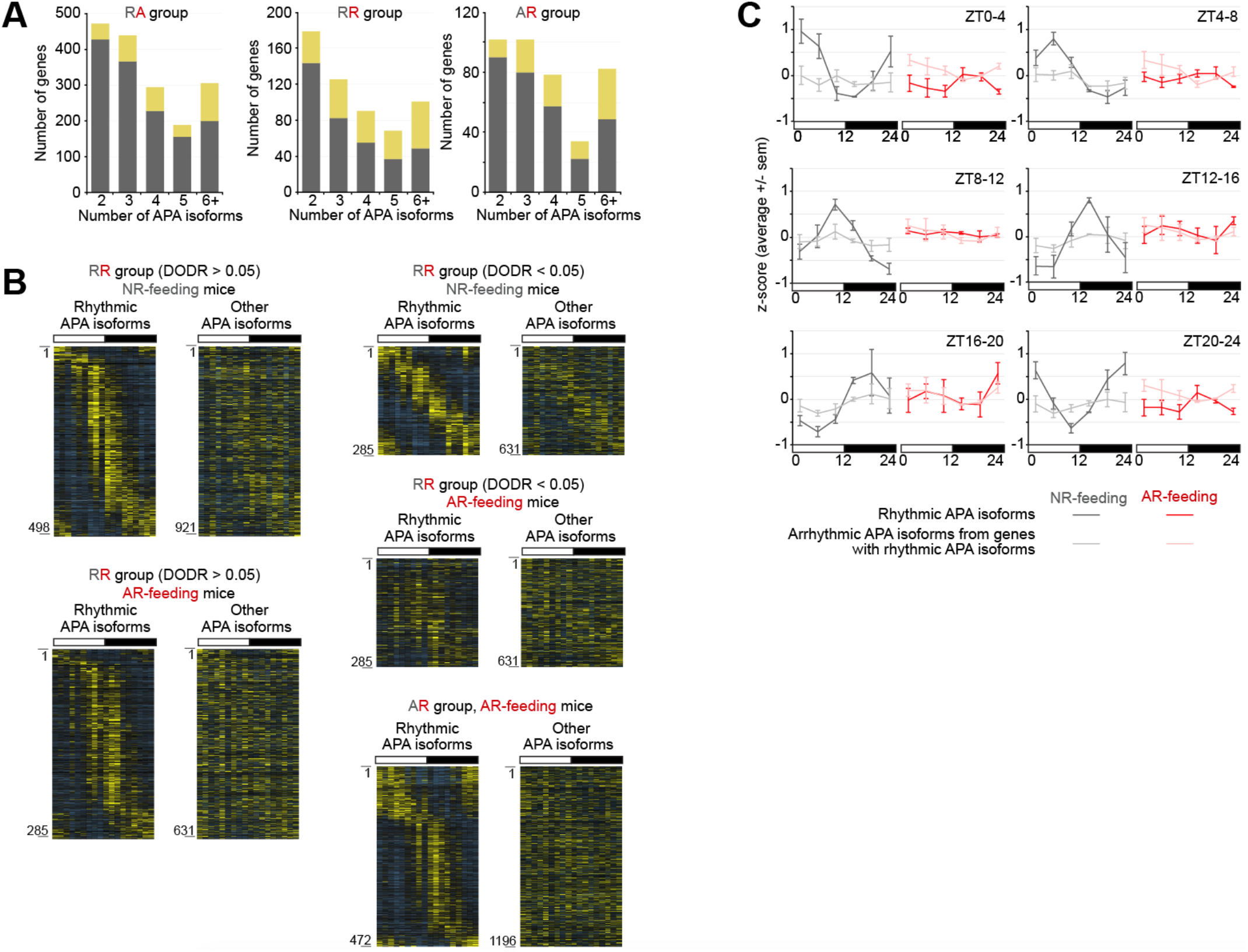
Analysis of APA isoform expression in the liver of mice fed only at night or fed arrhythmically. (related to Fig. 5) **(A)** Number of genes exhibiting differential rhythmicity for a single APA isoform or for multiple isoforms, in both AR-fed and NR-fed mice (RR group), NR-fed mice only (RA group), or AR-fed mice only (AR group). Number are parsed based on the number of APA isoforms per gene. **(B)** Heatmaps representing rhythmic APA isoforms (left) and arrhythmic APA isoforms located in a gene containing at least one rhythmic APA isoform (right). Heatmaps are shown for APA isoforms being rhythmically expressed in both AR-fed and NR-fed mice (RR group), NR-fed mice only (RA group), or AR-fed mice only (AR group). For the RR group, APA isoforms were further divided in two groups based on differential rhythmicity (DODR < 0.05). For the AR and RA groups, only genes showing differential rhythmicity (DODR, p<0.05) were considered. RNA-Seq signal for each isoform was mean-normalized, and the z-scores calculated for each gene based on all isoforms. For each heatmap pair, APA isoforms were ordered based on the phase of the rhythmic APA isoforms. Each heatmap column represent a single liver sample for AR- and NR-fed mice (n=18 total per condition; 6 timepoints, n=3 per time point). **(C)** Expression of APA isoforms being rhythmic in NR-fed mice only (RA group) (dark grey and dark red), and of arrhythmic APA isoforms transcribed from a gene harboring a rhythmic isoform in NR-fed mice only (light grey and pink). Expression corresponds to the average +/- sem of three mice per timepoint, and is parsed by bins of 4 hours based on peak phase of rhythmic APA isoform expression in NR-fed mice.

